# Fragmentation of Small-cell Lung Cancer Regulatory States in Heterotypic Microenvironments

**DOI:** 10.1101/2020.03.30.017210

**Authors:** Dylan L. Schaff, Shambhavi Singh, Kee-Beom Kim, Matthew D. Sutcliffe, Kwon-Sik Park, Kevin A. Janes

## Abstract

Small-cell lung cancers derive from pulmonary neuroendocrine cells, which have stemlike properties to reprogram into other cell types upon lung injury. It is difficult to uncouple transcriptional plasticity of these transformed cells from genetic changes that evolve in primary tumors or secondary metastases. Profiling of single cells also problematic if the required sample dissociation activates injury-like signaling and reprogramming. Here, we defined cell-state heterogeneities in situ through laser capture microdissection-based 10-cell transcriptomics coupled with stochastic-profiling fluctuation analysis. Using labeled cells from a small-cell lung cancer mouse model initiated by neuroendocrine deletion of *Rb1–Trp53*, we profiled variations in transcript abundance to identify cell-to-cell differences in regulatory state in vitro and in vivo. Fluctuating transcripts in spheroid culture were partly shared among *Rb1–Trp53*-null models, and heterogeneities increased considerably when cells were delivered intravenously to colonize the liver. Colonization of immunocompromised animals drove fractional appearance of alveolar type II-like markers and poised cells for paracrine stimulation from immune cells and hepatocytes. Immunocompetency further exaggerated the fragmentation of tumor states in the liver, yielding mixed stromal signatures evident in bulk sequencing from autochthonous tumors and metastases. Dozens of transcript heterogeneities recur irrespective of biological context; their mapped orthologs brought together observations of murine and human small-cell lung cancer. Candidate heterogeneities recurrent in the liver also stratified primary human tumors into discrete groups not readily explained by molecular subtype but with prognostic relevance. We conclude that heterotypic interactions in the liver and lung are an accelerant for intratumor heterogeneity in small-cell lung cancer.

**Statement of significance:** The single-cell regulatory heterogeneity of small-cell lung cancer becomes increasingly elaborate in the liver, a common metastatic site for the disease.

## Introduction

The categories, origins, and organization of tumor cell-to-cell heterogeneity are open questions of fundamental importance to cancer biology. Within normal tissues, single cells differ by lineage type and regulatory state. These distinctions blur, however, when cells lose their proper context because of tissue damage (1), transformation (2), or metastatic colonization (3). The details of such adaptive heterogeneity are expected to depend heavily on the originating cell type, the state of the cell when perturbed, and the local microenvironment where the cell resides.

Within the lung, the pulmonary neuroendocrine cell (PNEC) is a rare-but-important cell type that acts as an airway sensor for damaging stimuli (4). PNECs self-organize into 20–30-cell clusters at airway branch points through dynamic rearrangement of cell-cell contacts and reversible state changes suggesting epithelial-to-mesenchymal transition (EMT) (5). The latest evidence supports that certain PNECs have a reservoir of plasticity to convert into other lung cell types during tissue damage (1). Many tumors-metastases exhibit a state of chronic wounding (6), and PNECs are the main cell type of origin for small-cell lung cancer (SCLC) (7–9), a deadly form of lung carcinoma.

Regulatory mechanisms of SCLC plasticity are beginning to be dissected through systems-biology approaches (10,11) and genetically-engineered mouse models (GEMMs) (12). Human SCLC requires loss of *RB1* and *TP53* (13,14)—two tumor suppressors that also developmentally restrict pluripotency (15,16). GEMMs with *Rb1–Trp53* deleted by intratracheal delivery of Cre-expressing adenovirus (AdCMV-Cre) give rise to murine SCLCs similar to the classic ASCL1-high subtype of human SCLC (17,18). Deletion of additional tumor suppressors can synergize with *Rb1–Trp53* loss (19). For example, progression is accelerated by compound deletion of the Rb-family member *p130* (20). These GEMMs (7,8) and others (9) were instrumental in defining PNECs as the cell type of origin for SCLC.

Interestingly, phenotypes of the resulting murine tumors depend on the differentiation state of PNECs targeted for *Rb1–Trp53* deletion. Restricting adenoviral Cre to PNECs positive for the neuroendocrine marker Calca causes far fewer SCLCs to develop compared to when Cre expression is driven by a strong cytomegalovirus (CMV) promoter (21). Both tumor models are metastatic, but only the CMV-driven GEMM upregulates the transcription factor Nfib, which promotes widespread chromatin opening (22) and cell-lineage changes in both primary and metastatic sites (21). Many murine SCLC-derived cell lines are admixtures of cells with neuroendocrine and “non-NE” mesenchymal features (23). Other non-NE SCLC subpopulations are maintained by Notch signaling (12), which may also become activated in normal PNECs during injury-induced reprogramming (1). There might be other triggers of cellfate heterogeneity to uncover if SCLC regulatory states could be examined at single-cell resolution without injury-like dissociation of cellular context.

In this work, we examined the in situ transcriptomic regulatory heterogeneities of an established murine SCLC culture derived from an *Rb1^F/F^;Trp53^F/F^* animal administered AdCMV-Cre [KP1 cells (24); **Fig. 1A**]. Using GFP-labeled cells, fluorescence-guided laser capture microdissection (LCM), and 10-cell RNA sequencing (10cRNA-seq) (25), we considered three biological contexts: 1) tumor spheroids cultured in vitro and liver colonies in mice 2) lacking or 3) retaining an intact immune system (**Fig. 1B**). KP1 tumorspheres exhibited cell-to-cell regulatory heterogeneities in cell biology, aging, and metabolism that were shared with spheroid cultures of breast epithelia (25,26) and a separate SCLC line harboring additional tumorsuppressor deletions (19). Liver colonization gave rise to pronounced cell-state changes suggesting that paracrine signaling from the lung was partially resurrected in the liver. KP1 cells colonized in immunocompetent animals showed an exaggerated breadth of cell fates, with observed alveolar type II (ATII) markers intermingling with many non-NE stromal markers documented in SCLCs (23) and PNECs (1). Intersecting the three datasets yielded core recurrent heterogeneously expressed genes (RHEGs) and an in vivo RHEG set that was shared by all liver colonies but absent in tumorspheres. Core RHEGs from KP1 cells were broadly shared in bulk human SCLC transcriptomes, yet covariations among cases were not discriminating. By contrast, in vivo RHEGs showed weaker overall correlations but clustered human data into discrete prognostic groups that were separate from any 10-cell KP1 transcriptomes. The in vivo RHEGs defined here may reflect a set of injury-like SCLC adaptations that are possible during growth of tumors in the lung and metastases at different organ sites.

**Figure 1.**
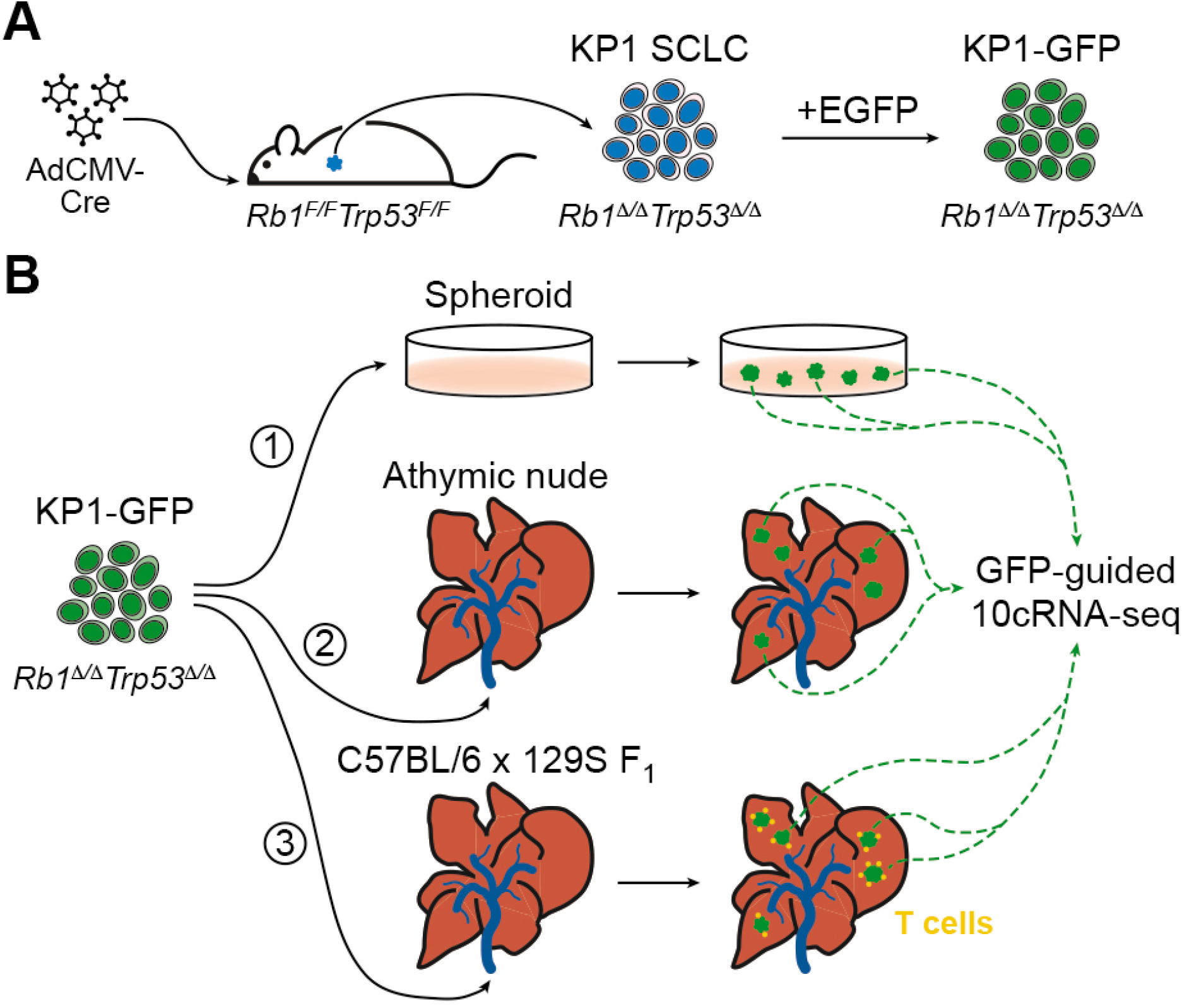
Stochastic profiling of transcriptional regulatory heterogeneity in three isogenic SCLC contexts. **A,** Derivation of KP1 small-cell lung cancer (SCLC) cells by intratracheal administration of adenovirus delivering cytomegalovirus promoter-driven Cre recombinase (AdCMV-Cre) to *Rb1^F/F^Trp53^F/F^* animals. The KP1 SCLC line was engineered to express ectopic enhanced green fluorescent protein (EGFP) for fluorescence-guided microdissection. **B,** The KP1-GFP derivative line was 1) cultured as three-dimensional spheroids in vitro or colonized to the liver of 2) athymic nude mice or 3) C57BL/6 x 129S F1 hybrid mice harboring an intact immune system. GFP-positive cells from multiple spheroids and liver colonies were randomly captured and measured by 10-cell RNA sequencing (10cRNA-seq) for stochastic profiling.

## Materials and Methods

### Cell and tissue sources

KP1 cells [generated by K.S.P. (24)] and *Chga-GFP;Rb*^Δ/Δ^;*p53*^Δ/Δ^;*p130*^Δ/+^;*Crebbp^-/-^*(RPC) cells [generated by K.B.K. (19)] were cultured as self-aggregating spheroids in RPMI medium 1640 (Gibco) with 10% FBS, 1% penicillin-streptomycin, and 1% glutamine. There was no cell-line authentication, and cells were not tested for mycoplasma contamination. KP1-GFP cells were prepared by transducing cells overnight with saturating lentivirus and 8 μg/ml polybrene as previously described (27). GFP-encoding lentivirus was prepared with pLX302 EGFP-V5 cloned by LR recombination of pLX302 (Addgene #25896) and pDONR221_EGFP (Addgene #25899). Stable transductants were selected with 2 μg/ml puromycin until control plates had cleared. Cultured KP1-GFP spheroids were kept to within 10 passages and cryoembedded as described previously (25).

To seed liver colonies, KP1-GFP spheroids were dissociated with 0.05% Trypsin/EDTA (Life Technologies), counted using a hemocytometer, and 2×10^5^ cells were injected via the tail vein of athymic nude (Envigo, *n* = 3 inoculations) or C57BL/6 x 129S F_1_ hybrid strain of mice (Jackson laboratory, *n* = 4 inoculations). Animals were not randomized. Liver colonies were resected after ~30 days and immediately cryoembedded in NEG-50, frozen in a dry ice-isopentane bath, and stored at −80°C (25). KP1 spheroids were cryosectioned at −24°C and liver colonies were cryosectioned at −20°C, both at 8 μm thickness as previously described (25). All mice were maintained according to practices prescribed by the National Institutes of Health in accordance with the IACUC protocol #9367. All animal procedures were approved by the Animal Care and Use Committee at the University of Virginia, accredited by the Association for the Assessment and Accreditation of Laboratory Animal Care (AAALAC).

### Fluorescence-guided LCM

Fresh frozen sections were fixed and dehydrated with ethanol and xylene as described previously for fluorescent cryosections (25). Freshly fixed samples were immediately microdissected on an Arcturus XT LCM instrument (Applied Biosystems) using Capsure HS caps (Arcturus). The smallest spot size on the instrument captured 3–5 SCLC cells per laser shot. 10cRNA-seq samples were obtained from 3–9 spheroids or liver colonies from one culture or animal, with independent follow-up by immunofluorescence (see below).

### RNA extraction and amplification

RNA extraction and amplification of microdissected samples was performed as described previously to minimize contaminating genomic amplification (25). Briefly, biotinylated cDNA was synthesized from RNA eluted from captured cells and purified with streptavidin magnetic beads (Pierce) on a 96 S Super Magnet Plate (Alpaqua). Residual RNA was degraded with RNAse H (NEB), and cDNA was poly(A) tailed with terminal transferase (Roche). Poly(A)-cDNA was amplified using AL1 primer (ATTGGATCCAGGCCGCTCTGGACAAAATATGAATTCTTTTTTTTTTTTTTTTTTTTTTTT) and a blend of Taq polymerase (NEB) and Phusion (NEB) for 25 cycles. RNA from bulk KP cell lines was isolated by RNEasy kit (Qiagen).

### 10-cell sample selection by quantitative PCR (qPCR)

Detection of transcripts by qPCR was performed on a CFX96 real-time PCR instrument (Bio-Rad) as described previously (28). 0.1 μl of preamplification material was used in the qPCR reaction. For each sample, we quantified the expression of *Gapdh* and *Rpl30* as loading controls. Samples were retained if the geometric mean quantification cycle of *Gapdh–Rpl30* was within 3.5x interquartile range of the median; samples outside that range were excluded because of over- or under-capture during LCM. For liver colonies, we also excluded samples with detectable quantification cycles of three high-abundance hepatocyte markers: *Alb, Fgb*, and *Cyp3a11.* qPCR primer sequences are available in Supplementary Table ST1.

### Library preparation

Ten-cell sequencing libraries were prepared by reamplification, purification, and tagmentation as described previously (25). Briefly, each poly(A) PCR cDNA sample was reamplified by PCR within its exponential phase (typically 10 to 20 cycles). Re-amplified cDNA was then twice purified with Ampure Agencourt XP SPRI beads, and samples were quantified on a CFX96 real-time PCR instrument (Bio-Rad) using a Qubit BR Assay Kit (Thermo Fisher). Samples were diluted to 0.2 ng/μl and tagmented with the Nextera XT DNA Library Preparation Kit (Illumina). Bulk KP libraries were prepared from 500 ng of total RNA by the Genome Analysis and Technology Core at the University of Virginia using mRNA oligo dT-purified with the NEB Next Ultra RNA library preparation kit (NEB).

### RNA sequencing

10cRNA-seq data were sequenced and aligned as previously described (25). Ten-cell samples were multiplexed at an equimolar ratio, and 1.3 pM of the multiplexed pool was sequenced on a NextSeq 500 instrument with NextSeq 500/550 Mid/high Output v1/v2/v2.5 kits (Illumina) to obtain 75-bp paired-end reads. Bulk KP RNA samples were sequenced on a NextSeq 500 to obtain 50-bp single-end reads. Adapters were trimmed using fastq-mcf in the EAutils package (version ea-utils.1.1.2-779) with the following options: -q 10 -t 0.01 -k 0 (quality threshold 10, 0.01% occurrence frequency, no nucleotide skew causing cycle removal). Quality checks were performed with FastQC (version 0.11.8) and multiqc (version 1.7). Mouse datasets were aligned to the mouse transcriptome (GRCm38.82), reference sequences for ERCC spikeins, and pLX302-EGFP by using RSEM (version 1.3.0) and Bowtie 2 (version 2.3.4.3). RSEM processing of the 10cRNA-seq data also included the following options: --single-cell-prior --paired-end. Counts from RSEM processing were converted to transcripts per million (TPM) by dividing each value by the total read count for each sample and multiplying by 10^6^. Total read count for TPM normalization did not include mitochondrial genes or ERCC spike-ins.

### Whole-exome sequencing and identification of KP1 polymorphisms

Genomic DNA was prepared from KP1 spheroid cultures with the DNeasy Blood & Tissue kit (Qiagen). Whole-exome sequencing at 100x coverage was performed as a contract service with Genewiz as described in the accompanying contribution (29) but aligning to the mouse reference GRCm38 to generate the VCF for downstream analysis. Distinguishing polymorphisms were defined as homozygous variants in the 3’ untranslated region of transcripts with a median 10cRNA-seq abundance greater than 100 TPM.

### RNA FISH

A 150-bp fragment of human *SOX4* was cloned into pcDNA3, used as a template for in vitro transcription of a digoxigenin-labeled riboprobe for RNA FISH, and imaged as previously described (26). Loading-control riboprobes for *GAPDH, HINT1*, and *PRDX6* were previously reported (27).

### Immunohistochemistry

Staining was performed by the Biorepository and Tissue Research Facility at the University of Virginia with 4 μm paraffin sections. For F4/80, antigen retrieval and deparaffinization were performed in PT Link (Dako) using low pH EnVision FLEX Target Retrieval Solution (Dako) for 20 minutes at 97°C. Staining was performed on a robotic platform (Autostainer, Dako). Endogenous peroxidases were blocked with peroxidase and alkaline phosphatase blocking reagent (Dako) before incubating the sections with F4/80 antibody (AbD Serotech, #MCA497R) at 1:200 dilution for 60 minutes at room temperature. Antigen–antibody complex was detected by using rabbit anti-rat biotin and streptavidin-HRP (Vector Laboratories) followed by incubation with 3,3’-diaminobenzidine tetrahydrochloride (DAB+) chromogen (Dako). For CD3, sections were deparaffinized using EZ Prep solution (Ventana), and staining was performed on a robotic platform (Ventana Discover Ultra Staining Module). A heat-induced antigen retrieval protocol set for 64 min was carried out using Cell Conditioner 1 (Ventana). Endogenous peroxidases were blocked with peroxidase inhibitor (CM1) for 8 minutes before incubating the section with CD3 antibody (Dako, #A0452) at 1:300 dilution for 60 minutes at room temperature. Antigen-antibody complex was detected using DISCOVERY OmniMap antirabbit multimer RUO detection system and DISCOVERY ChromoMap DAB Kit (Ventana). All slides were counterstained with hematoxylin, dehydrated, cleared, and mounted for assessment. For both F4/80 and CD3 stains, cells were counted visually and reported as the average of multiple 10x-field images surrounding individual KP1 cell colonies in liver sections obtained from athymic nude and C57BL/6 x 129S F1 hybrid mice.

### Immunofluorescence and image segmentation

For KP1 spheroids, cultures were fixed, embedded, and cryosectioned as described previously for MCF10A-5E spheroids (27). The immunostaining procedure was very similar but substituted the basic M.O.M. immunodetection kit (Vector Laboratories) as the blocking step and M.O.M. diluent for incubation with primary antibodies overnight at room temperature. Primary antibodies recognizing the following proteins were used: GFP (Abcam #ab13970, 1:500), Kpna2 (Thermo Fisher #MA5-31790, 1:500), Cse1l (Abcam #ab96755, 1:500), Mki67 (Thermo Fisher #MA5-14520, 1:500), Rps6 (Cell Signaling #2317, 1:50), ubiquitin (Santa Cruz #sc-8017, 1:200), Mavs (Cell Signaling #4983, 1:50), Hif1α (R&D Systems #NB100-134, 1:200). Images were collected on >15 spheroids from two separately-stained pairs of cryosections.

For KP1 liver colonies, formalin-fixed and paraffin embedded tissue sections were processed and immunostained as described previously (30). Primary antibodies recognizing the following proteins were used: GFP (Abcam #ab13970, 1:500), Cd74 (BD Biosciences #555317, 1:200). The GFP antibody was paired with Alexa 647-conjugated anti-chicken secondary antibody (Invitrogen) to avoid confounding tissue autofluorescence. Images were collected on 5–12 liver colonies from 2–4 animals.

Image segmentation and single-cell analysis of immunoreactivity was based off of an earlier pipeline (30). For Kpna2–Cse1l coregulation, DAPI-positive nuclei were smoothed with a Gaussian filter and segmented by global two-class thresholding with Otsu’s method. The borders of each cell were identified using the “propagation” method based on GFP expression in the Alexa Fluor 488 channel. Median object intensities were measured in the Alexa Fluor 488, Alexa Fluor 555, and Alexa Fluor 647 channels for analysis. For core and periphery measurements (Mki67, Mavs, Rps6, Ub, and Hif1α), borders of entire spheroids were manually circled and then shrunk by 50 pixels (16 μm). Cells within the shrunk boundary were identified as core cells while cells in the tumorsphere but outside of the core boundary were identified as periphery cells. Cells within 75 pixels (24 μm) of the image boundary were eliminated to avoid ambiguity of core or periphery determination. Core and periphery median object intensities were separately aggregated in R, normalized to the overall median intensity for the region of the spheroid, and fit to a lognormal distribution or a mixture of lognormal distributions with unequal variance. Mavs^+^ spots of 2–10 pixels (0.64–3.2 μm) diameter were smoothed with a Gaussian filter, segmented by global two-class thresholding with Otsu’s method at a minimum threshold of 0.0167, and related to the cell containing the spot. For KP1 liver colonies, DAPI-positive nuclei were smoothed with a Gaussian filter and segmented by global two-class thresholding with Otsu’s method. Nuclei were dilated by 5 pixels (1.6 μm) to capture the vicinity of each cell, and median object intensities were measured in the Alexa Fluor 555 and Alexa Fluor 647 channels for analysis. Median object intensities were aggregated in R, normalized to the overall median intensity, and fit to a lognormal distribution or a mixture of lognormal distributions with unequal variance. Normalized median intensity gates for positive cells were set at the 99^th^ percentile of the leftmost distribution.

### Cell dissociation

KP1 spheroids were centrifuged, washed in PBS, and treated with 0.05% trypsin-EDTA (Gibco) or 1x accutase (Invitrogen) for five minutes at 37°C followed by lysis for protein analysis as previously described (31). For RNA analysis of Notch target genes, the dissociation was stopped with growth medium and cells were pelleted then washed in PBS. After centrifugation, cells were resuspended in growth medium for one hour before lysis and RNA purification with the RNeasy Mini Plus Kit (Qiagen). Control cells were treated identically except for the five-minute addition of dissociating enzyme.

### Immunoblot analysis

Quantitative immunoblotting was performed as previously described (31). Primary antibodies recognizing the following proteins were used: Notch2 (Cell Signaling #5732, 1:1000), vinculin (Millipore #05-386, 1:10,000), GAPDH (Ambion #AM4300, 1:20,000), tubulin (Abcam #ab89984, 1:20,000).

### Mouse-to-human ortholog mapping

Human orthologs for mouse genes were obtained from the Ensembl biomart in R using the getAttributes function. For genes with multiple human-ortholog mappings, we used expression characteristics of the human datasets considered [MCF10A-5E (25) and human SCLC (32)] to favor more-reliable clustering afterwards. For mouse genes with two human mappings, the human ortholog with higher expression variance in the corresponding human dataset was retained. For mouse genes with greater than two human mappings, two orthologs with the highest expression correlation were identified. From these, the ortholog with the higher expression variance was retained, as in the two-mapping case. Any remaining mouse gene names were capitalized in accordance with human gene symbol conventions.

### Overdispersion-based stochastic profiling

Stochastic profiling with 10cRNA-seq data was performed exactly as described in the accompanying contribution (29). For RPC spheroids, the adjusted variance of KP1 spheroid split-and-pool controls was used as the threshold for candidate heterogeneities.

### Robust identification of transcriptional heterogeneities through subsampling

To minimize the contribution of outliers to the overdispersion analysis in samples collected from liver colonies, we generated 100 subsampled simulations for overdispersion-based stochastic profiling. After sample selection (see above), there were 33 10-cell samples plus 35 pool-and-split controls for nude liver colonies and 31 10-cell samples plus 24 pool-and-split controls for C57BL/6 x 129S F1 hybrid liver colonies. For each dataset, overdispersion-based stochastic profiling was performed 100 times with random downsampling to 28 10-cell samples and 20 pool-and-split controls (29). Only genes that recurred as candidates in >75% of simulations were evaluated further as candidate heterogeneously expressed genes.

### Filtering out hepatocyte contamination in heterogeneous expressed genes

Among overdispersed transcripts in liver colonies, we further excluded genes that might vary because of residual hepatocyte capture during LCM. For each 10-cell sample, we calculated the geometric mean abundance of 11 liver-specific markers (*Alb, Fgb, Cyp3a11, Ambp, Apoh, Hamp, Ass1, Cyp2f2, Glul, Hal*, and *Pck1*) from published studies (33–36). Candidates that were significantly correlated with the mean liver signature (*p* < 0.05 by Fisher Z-transformed Spearman *ρ* correlation) were removed from further consideration for the in vivo study.

### Continuous overdispersion analysis

Overdispersion values from the 2007 transcripts identified as candidate heterogeneities in either the KP1 spheres or in vivo conditions were recorded for 100 subsampling iterations. For the KP1 in vitro spheroids, 100 iterations of leave-one-out crossvalidation were performed as detailed in the accompanying contribution (37). Transcripts were retained if the 5th percentile of overdispersion in the C57BL/6 x 129S F1 hybrid condition was greater than the 95th percentiles of the other two conditions. If a gene was not expressed in a condition, the 5th and 95th percentiles were set to zero, and the gene was assigned to the overall median overdispersion during clustering.

### Statistics

Sample sizes for stochastic profiling were determined by Monte Carlo simulation (38). Significance of overlap between candidate genes in KP1 spheroids and MCF10A-5E spheroids was evaluated using the hypergeometric test using the “phyper” function in R and a background of 20,000 genes. Pearson correlation between pairs of transcripts detected in both KP1 and MCF10A-5E spheroids were assessed using the “cor.test” function. Significant increases in number of candidate genes between different conditions were assessed by the binomial test using the “binom.test” function in R. Spearman *ρ* correlation between overdispersed transcripts and liver markers was calculated using the “cor.test” function. Spearman *ρ* correlations were Fisher Z-transformed using the “FisherZ” function from the R package “DescTools” (version 0.99.31). Co-occurrence of transcript fluctuations was evaluated by hypergeometric test after binning 10cRNA-seq above or below the geometric mean of the two transcripts compared (1 – *p* evaluated counter-occurrence). Differences in cell number by immunohistochemistry were assessed by the Wilcoxon rank sum test using “wilcox.test”. Significance of overlaps between candidate genes identified in spheroids, nude mice, and C57BL/6 x 129S F1 hybrid mice were assessed by Monte Carlo simulations and corrected for multiple hypothesis testing as described (29). Significance of differences in protein abundance by immunoblotting were assessed by one-way ANOVA with Tukey HSD post-hoc test. Differences in log-transformed mRNA abundance by qPCR were assessed by Student’s *t* test with Šidàk correction for multiple comparisons. Hierarchical clustering was performed using “pheatmap” with standardized values, Euclidean distance, and “ward.D2” linkage or non-standardized values, Pearson distance, and “ward.D2” linkage. Consensus clustering of standardized values was performed with ConsensusClusterPlus (version 1.48.0) (39) with a range of 1–10 consensus clusters, subsampling of both 80% the input samples and 80% of the genes within each sample, 1000 iterations, Euclidean distance, and “ward.D2” linkage. Gene set enrichment analyses were performed through the Molecular Signatures Database (40). Overlaps between gene lists and hallmark gene sets were computed using a hypergeometric test with false-discovery rate correction for multiple comparisons.

### Data availability

Bulk and 10cRNA-seq data from this study is available through the NCBI Gene Expression Omnibus (GSE147358, https://www.ncbi.nlm.nih.gov/geo/query/acc.cgi?acc=GSE147358 Reviewer token: ufczqoiwzbwjnet). Other RNA-seq datasets were obtained from the Gene Expression Omnibus (GEO): MCF10A-5E 10cRNA-seq (GSE120261), AdCMV-Cre and AdCalca-Cre GEMM (GSE116977), and human SCLC tumor (GSE60052).

## Results

### Study design and rationale

We sought to define how SCLC regulatory heterogeneity was compiled in different microenvironments. To avoid spurious variations among GEMM tumors that arise autochthonously, we used KP1 cells, a polyclonal *Trp53*^Δ/Δ^*Rb1*^Δ/Δ^ line derived from a tumor initiated by intratracheal administration of AdCMV-Cre. We sequenced the bulk transcriptome of KP1 cells and found that they were very similar to three other *Trp53*^Δ/Δ^*Rb1*^Δ/Δ^ lines prepared in similar genetic backgrounds (GSE147358). By contrast, autochthonous SCLC tumors from related GEMMs (21) were different and also more variable among primary tumors, as expected (Supplementary Fig. S1). Before starting, we genetically labeled KP1 cells with EGFP for unambiguous isolation of cells administered in vivo (**Fig. 1A**).

SCLCs frequently metastasize to the liver (41). We mimicked the terminal steps of metastatic colonization and outgrowth by tail-vein injection of EGFP-labeled KP1 cells, which readily establish lesions in the livers of athymic nude mice (**Fig. 1B** and Supplementary Fig. S2A). Although KP1 cells have a mixed genetic background, we discovered that subcutaneous allografts and liver colonies were 100% successful in first-generation crosses (F1) of C57BL/6 and 129S inbred strains (Supplementary Fig. S2B). Inoculating C57BL/6 x 129S F1 hybrid animals thus afforded a third setting in which liver colonization and expansion could occur in the presence of a cell-mediated immune response.

### KP1 tumorspheres share adaptive transcriptional regulatory heterogeneities with breast-epithelial spheroids and other SCLC tumorspheres

Cultured KP1 cells grow as spheroidal aggregates in which cell-cell interactions may contribute to the overall heterogeneity of the population (12). To preserve these interactions, we processed KP1 spheroids by cryoembedding within seconds and sectioning–staining exactly as if they were tissue (25). Cells were microdissected from the outermost periphery of each spheroid to ensure that all cells profiled had equal availability of nutrients. We gathered 10-cell pools across multiple spheroids to average out subclonal differences within the line and highlight pervasive heterogeneities that characterize spheroid culture. Using 10cRNA-seq (25), we measured the transcriptomes of 28 separate 10-cell groups of KP1 cells along with 20 pool-and-split controls as 10-cell equivalents obtained by LCM. The data were analyzed for candidate regulatory heterogeneities by stochastic profiling (26) implemented with an overdispersion metric optimized for RNA-seq, as described in the accompanying contribution (29). The analysis yielded 405 candidate genes that were much more variable in the 10-cell samples than expected given their average abundance and technical reproducibility (**Fig. 2A** and Supplementary File S1).

**Figure 2.**
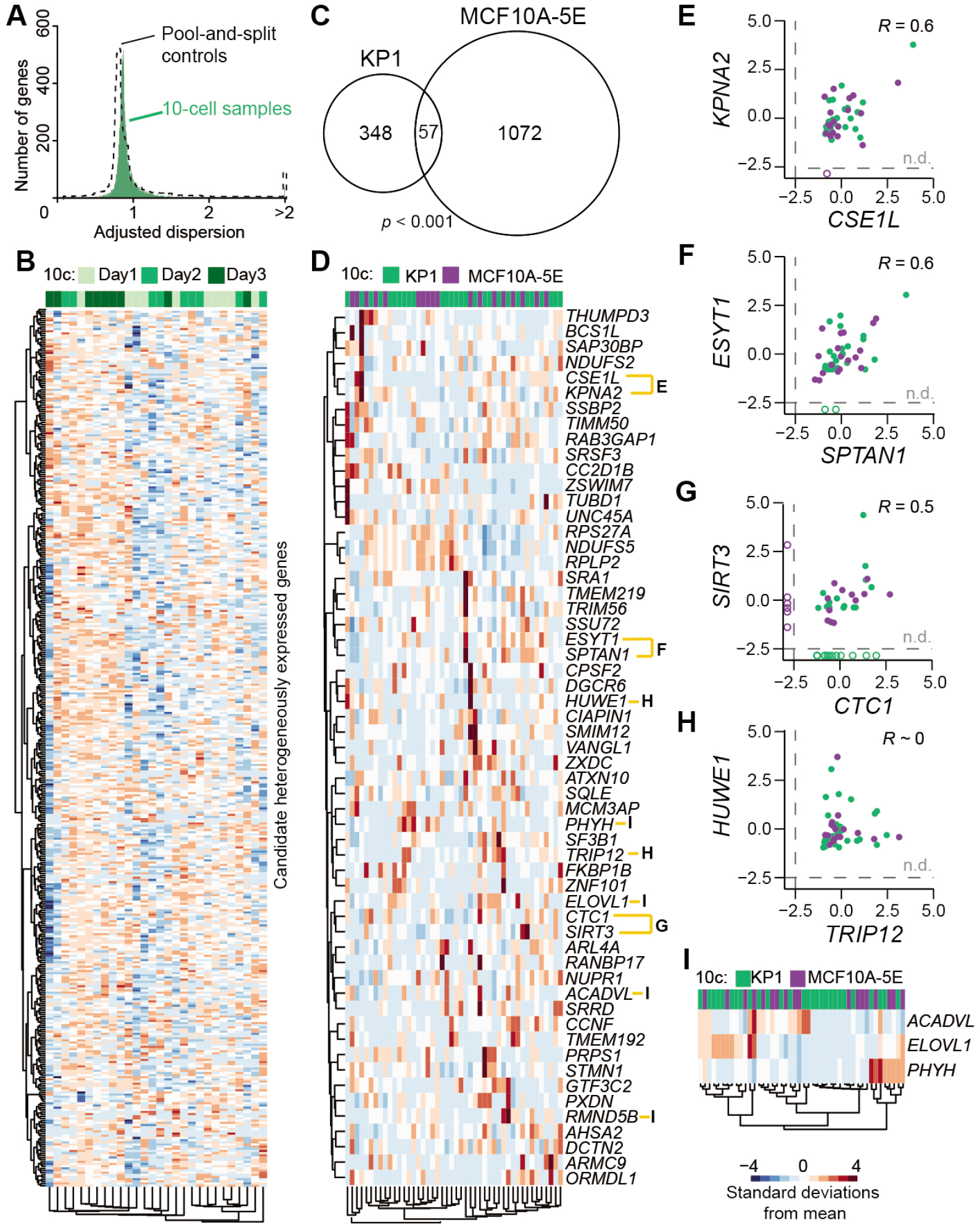
Shared transcriptional regulatory heterogeneities between KP1 spheroids and MCF10A-5E breast-epithelial spheroids. **A,** Overdispersion plot showing 10-cell sample distribution for KP1 spheroids (green) overlaid on pool-and-split controls of 10-cell equivalents (dashed). **B,** Clustergram of the 405 transcripts identified as candidate heterogeneities within KP1 spheroids. Sample acquisition days are annotated. Data were log transformed before standardization. **C,** Venn diagram of orthologous candidates between KP1 spheroids and MCF10A-5E spheroids analyzed in Supplementary Fig. S3. Significance of the intersection was assessed by hypergeometric test with 12,612 total detectable transcripts in KP1 cells and 12,927 total detectable transcripts in MCF10A-5E cells. **D,** Clustergram of the spheroid RHEGs annotated by human ortholog. **E–H,** Pairwise Pearson correlations between the indicated gene pairs in **D** among samples where both genes were detected (filled). n.d., not detected. **I,** Clustergram of three transcripts encoding enzymes for lipid metabolism. In **D–I**, 10-cell samples of KP1 cells (green) and MCF10A-5E cells (purple) were standardized separately by z-score before clustering or correlation.

Samples were collected across multiple days to assess whether batch effects dominated the fluctuation analysis. We clustered gene candidates hierarchically and asked whether the fluctuation signatures clustered according to when the 10-cell samples were collected (**Fig. 2B**). Each grouping was comprised of 10-cell profiles from all batches, supporting that the analytical strategy was robust amidst day-to-day variations in LCM, RNA extraction, and sample amplification. Standard gene set enrichment analysis indicated hallmarks for cell-cycle transitions, Myc–mTORC1 signaling, and metabolism (Supplementary File S2), consistent with the variable growth of spheres in the culture.

Previously, our group used 10cRNA-seq to revisit an earlier analysis of transcriptional regulatory heterogeneity in 3D cultured MCF10A-5E breast-epithelial spheroids (25,26). With an analytical pipeline for stochastic profiling by 10cRNA-seq now in hand (29), we quantified the gene-by-gene overdispersion and identified 1129 candidate heterogeneities (Supplementary Fig. S3A and S3B). The list included multiple transcripts that were independently validated to be heterogeneous by RNA fluorescence in situ hybridization (26,42), including one transcript (*SOX4*) that we validated here (Supplementary Fig. S3B and S3C). The analysis provided a second context for regulatory heterogeneity that exists during spheroidal growth.

Murine SCLC cells and human breast epithelial cells are undoubtedly very different, but normal PNECs derive from an epithelial lineage (5) and often adopt a columnar morphology similar to that seen in the breast. The KP1 study detected significantly fewer genes as regulated heterogeneously compared to MCF10A-5E (*p* < 10^-15^ by binomial test), corroborating the differences in spheroid culture format. MCF10A-5E cells were 3D cultured in reconstituted basement membrane, which traps secreted factors locally around the spheroids (31), whereas KP1 spheroids develop freely in suspension. Despite differences in the overall number of candidates, we found significant overlap in shared genes after mapping mouse and human orthologs (see Materials and Methods; **Fig. 2C**). Intersecting the two gene groups only marginally enriched for cell-cycling transcripts (six of 57 genes, *p* = 0.04 by hypergeometric test). This result corroborates findings of the accompanying work (29) that cycling transcripts do not strongly confound stochastic profiling experiments. The intersected spheroid data suggested other biological processes in addition to proliferation and raised the possibility that cell growth–competition within epithelial spheroids elicits a set of RHEGs, which generalize beyond a specific culture format.

We next asked whether there might be any common heterogeneities in regulation between the two contexts after correcting for transcript abundance. When the standardized fluctuations of KP1 and MCF10A-5E spheroids were coclustered by gene ortholog, there were multiple close pairings consistent with shared biology or biological category (**Fig. 2D**). For instance, the importin *KPNA2* covaried with its exportin, *CSE1L* (**Fig. 2E**) (43). Single-cell Kpna2–Cse1l coregulation was independently confirmed by immunofluorescence of cryosectioned KP1 spheroids (Supplementary Fig. S4). We also observed cross-species correlations in genes functioning at the interface of the plasma membrane and endoplasmic reticulum: *ESYT1* and *SPTAN1* (**Fig. 2F**) (44). Although near the detection limit for both cell types, we noted cofluctuations in *SIRT3* and *CTC1*, two factors implicated in cellular longevity (**Fig. 2G**) (45,46). Together, these gene pairings provide starting points for unraveling singlecell regulatory pathways that become co-activated when epithelia proliferate outside of their normal polarized context.

Elsewhere among the spheroid RHEGs, we found instances of mutually exclusive transcript heterogeneities, such as with *HUWE1* and *TRIP12* (**Fig. 2H**). Although these E3 ubiquitin ligases have been separately tied to cancer, they converge as independent triggers of ubiquitin fusion degradation (47), an unusual proteasomal pathway not studied in cancer. Separately, we recognized a preponderance of metabolic enzymes related to lipids and clustered the fatty acid elongase *ELOVL1*, the β-oxidation dehydrogenase *ACADVL*, and the α-oxidation hydroxylase *PHYH* (**Fig. 2I**). Even with 10-cell pooling, we rarely observed these enzymes abundantly expressed in the same sample, suggesting independent states of lipid synthesis and degradation that could be mined deeply in the future for covariates. Expanding candidate lists around positive and negative covariates has proved powerful in mechanistic follow-on work (30,31).

A more-focused question is whether the regulatory heterogeneities of KP1 spheroids generalize to other SCLC contexts. We addressed this by profiling spheroid cultures of precancerous *Chga-GFP;Rb^Δ/Δ^;p53^Δ/Δ^;p130^Δ/+^* cells transformed by compound deletion of the SCLC tumor suppressor, *Crebbp* [RPC cells, (19)]. Following the same study design as KP1 spheroids, stochastic profiling by 10cRNA-seq identified 817 genes as candidates for heterogeneous regulation (Supplementary File S1). The ~twofold increase in gene number compared to KP1 spheroids (*p* < 10^-36^ by binomial test) is consistent with the increased EMT-like plasticity conferred by loss of *Crebbp* (19). In addition to the overall increase in heterogeneity, RPC spheroids shared 79 candidates in common with KP1 spheroids (Supplementary Fig. S5A). The overlap was highly significant (*p* < 10^-20^ by hypergeometric test) and included the previously described *Kpna2, Cse1l*, and *Phyh* transcripts that recurred in MCF10A-5E spheroids (Supplementary Fig. S5B). The KP1–RPC intersection also contained multiple kinases (*Pbk, Prkar2a*), phosphatases (*Ppp1cc, Ppp2r1a, Ppp6c*), epigenetic regulators (*Hdac2, Sin3a*), and nuclear receptor effectors (*Ncor1, Ncoa4).* Results from the two lines profiled suggest that SCLC cells variably reconfigure their signaling, transcriptional, and epigenetic states, even in monoculture.

### SCLC reprogramming and paracrine signaling are initiated by colonization of KP1 cells to the liver

To begin examining how heterotypic interactions augment SCLC regulatory heterogeneity, we dissociated KP1 spheroids and colonized the liver of athymic nude mice (**Fig. 1B**). We ensured that SCLC-hepatocyte communication was reflected in the 10cRNA-seq data by sampling the margins of separate GFP^+^ KP1 liver colonies, analogous to the spheroid margins sampled in vitro (**Fig. 3A**). Focusing on the KP1–hepatocyte interface implied that some level of cell contamination would be introduced by collateral pickup during the LCM step. We rigorously controlled for hepatocyte contamination through a two-step negative-selection procedure. Samples were excluded if hepatocyte markers were abundant by qPCR (Supplementary Table ST1), and transcripts were removed post-analysis if they covaried with the residual hepatocyte content in the sequenced sample (**Fig. 3B**). Additionally, we oversampled the in vivo samples, collecting 33 10-cell pools that were subsampled 100 times as random groups of 28 for the dispersion analysis (see Materials and Methods; **Fig. 3B** and Supplementary Fig. S6). The pipeline collectively identified 898 robust candidates fluctuating independently of residual liver markers and appearing in ≥75% of subsampling runs (**Fig. 3C** and Supplementary File S1).

**Figure 3.**
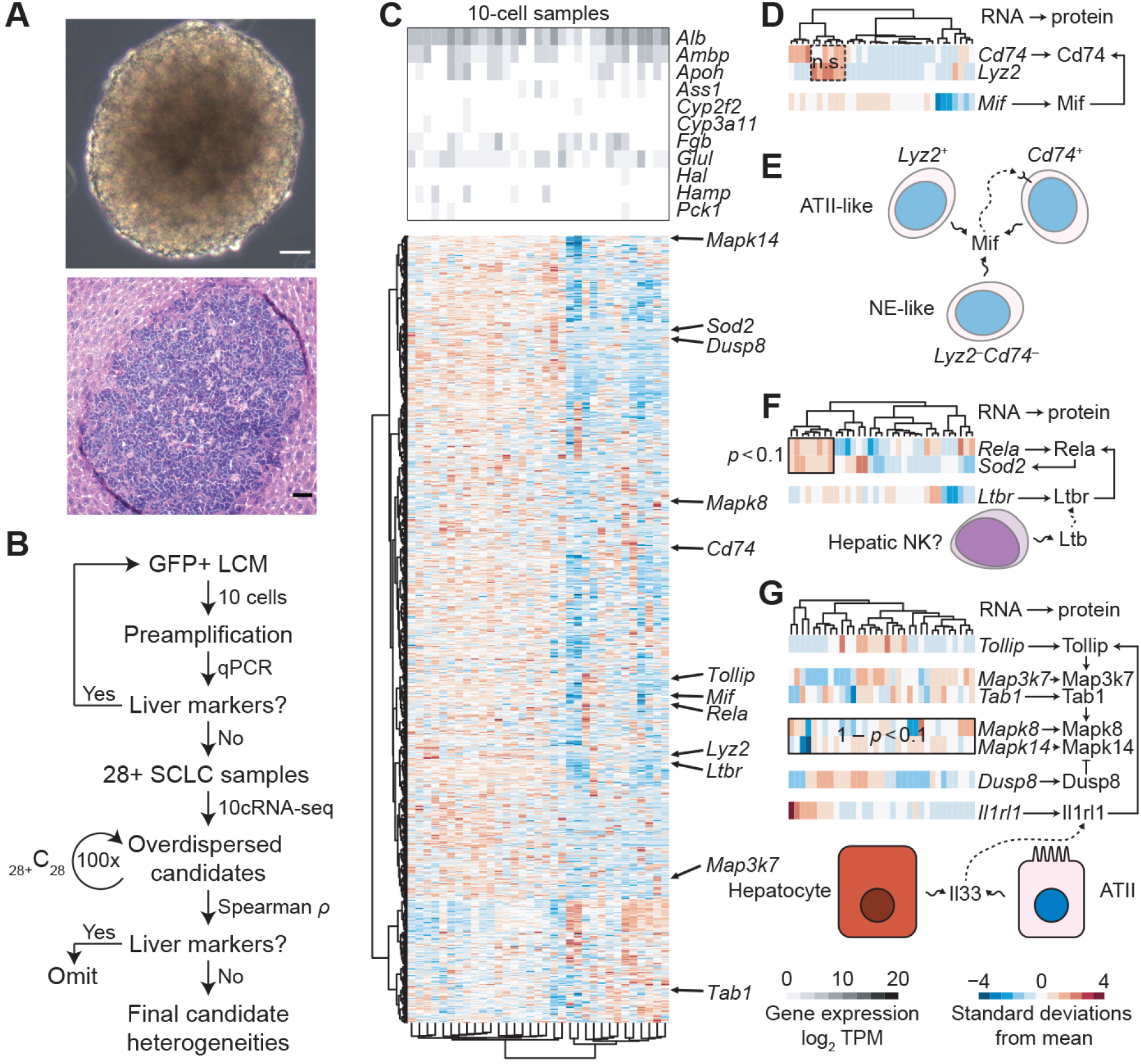
KP1 liver colonization in athymic nude mice causes partial reprogramming and engages heterotypic paracrine-signaling networks. **A,** Phase-contrast image of cultured KP1 spheroids (upper) compared to a brightfield hematoxylin-eosin stain of a KP1 liver colony in an athymic nude mouse (lower). Scale bar is 80 μm. **B,** Flowchart illustrating the experimental and analytical strategy controlling for liver contamination in 10cRNA-seq data and in candidate heterogeneities identified by stochastic profiling. Subsampling results from the 100 dispersion analyses of 28 10-cell samples are shown in Supplementary Fig. S6. **C,** Relative abundance of liver markers (upper) and log-standardized 10-cell sampling fluctuations of robust candidate heterogeneities identified by stochastic profiling (lower). **D,** Log-standardized sampling fluctuations of the alveolar type II (ATII) markers *Cd74* and *Lyz2* together with the Cd74 ligand, *Mif.* **E,** Schematic illustrating the hypothesized relationship between neuroendocrine (NE)- and ATII-like states and Mif signaling. **F,** Log-standardized sampling fluctuations of the *Rela* transcription factor, *Sod2* (a Rela target gene), and *Ltbr* (a Rela-inducing receptor). The ligand for Ltbr is produced by liver-resident NK cells (33). **G,** Log-standardized sampling fluctuations of the indicated signaling transcripts and their pathway relationships downstream of the Il33 receptor, Il1rl1, which is present in KP1 cells but not heterogeneously regulated. Il33 is produced by hepatocytes in the liver and ATII cells in the lung. For **D, F,** and **G,** enriched or unenriched coexpression was evaluated by hypergeometric test of 10-cell observations above their respective log means. n.s., *p* > 0.1 and 1 – *p* > 0.1.

With the 10cRNA-seq data in hand, we addressed two additional concerns about the allograft sampling. First, dissociation of KP1 spheroids before inoculation may have disrupted spatial gradients in gene expression and led to clonal sub-selection of non-peripheral cells in vivo. We sought out such possibilities by looking for heterogeneous candidates in spheroids, which were not overdispersed in the liver. For two such candidates—Mki67 and Mavs—we confirmed by immunofluorescence that their expression was comprised of multiple states in the spheroid core and at the periphery (Supplementary Fig. S7A and S7B). In contrast, two controls that were not overdispersed either in vitro or in vivo (Rps6 and Ubc) were captured by a single-state model in the periphery and showed limited spatially segregated expression in the spheroid core (Supplementary Fig. S7C and S7D). Thus, clonal sub-selection of KP1-spheroid niches is not pervasive in KP1 liver colonies.

Second, 10cRNA-seq fluctuations in vivo could have arisen from contaminating cells other than hepatocytes. We addressed this issue by whole-exome sequencing of KP1 cells to identify distinguishing single-nucleotide polymorphisms (SNPs). For homozygous variants reliably detected by 10cRNA-seq, we consistently found >99% allelic concordance between bulk RNA-seq of cultured KP1 cells and 10cRNA-seq from liver colonies (Supplementary Fig. S8A). Similar purity was observed for a homozygous pair of SNPs in linkage disequilibrium and absent from both the C57BL/6 and 129S genomes (Supplementary Fig. S8B). We conclude that 10cRNA-seq transcriptomes in this study are overwhelmingly KP1 in origin.

Enriched gene sets for in vivo candidates were virtually identical to spheroid cultures, except for the addition of STAT5 and interferon γ hallmarks likely resulting from innate immune responses (Supplementary File S2). Beyond the significant increase in candidates (*p* < 10^-15^ by binomial test), we noted that sample-to-sample fluctuations were qualitatively more dramatic on the margin of liver colonies when compared to KP1 spheroids (**Fig. 2B** and **3C**). Multiple, smaller subsets of candidates were especially interesting. For example, among the robust candidates were the alveolar type II (ATII) markers *Cd74* (48) and *Lyz2* (49), as well as the Cd74 ligand, *Mif.* Surprisingly, when the sample-by-sample fluctuations of these three genes were clustered, we did not detect any significant co-occurrence that would have suggested full transdifferentiation to an ATII phenotype (see Materials and Methods; **Fig. 3D**). The results agree with scRNA-seq data obtained in deprogrammed PNECs (1), where *Cd74* and *Lyz2* markers are anti-correlated among cells with a non-NE phenotype (Supplemental Fig. S9). The patterns detected by stochastic profiling suggest that a subset of KP1 cells reprogram into partial ATII-like states, only one of which senses Mif produced locally (**Fig. 3E)**.

Other groups of transcripts required inputs from non-KP1-derived cell types in the liver to rationalize. Two robust candidates were the NF-κB subunit *Rela* and an NF-κB target gene, *Sod2* (50), which co-occurred strongly (*p* < 0.1) when considering that NF-κB is mostly regulated posttranslationally (**Fig. 3F**). Within the candidate heterogeneities, we also identified the NF-κB-inducing receptor *Ltbr*(51), which varied separately from *Rela–Sod2.* However, the Ltbr ligand (*Ltb*) was effectively absent in KP1 cells (less than 1.5 TPM in bulk samples and pool-and-split controls from liver colonies). We searched Tabula Muris (33) and found that *Ltb* is abundantly expressed in hepatic natural killer (NK) cells, the most-prevalent lymphocyte population in the liver. Given that NK cell activity is retained or enhanced in athymic nude mice (52), their paracrine communication with KP1 cells is a plausible mechanism for heterogeneous NF-κB pathway activation in the liver.

We also found evidence for variable regulation in signal transducers of interleukin 1-family cytokines. The inhibitory adaptor *Tollip* (53), the mitogen-activated protein kinase (MAPK) kinase kinase *Map3k7* and its activator *Tab1* (54), the downstream stress-activated MAPKs *Mapk8* (or *Jnk1*) and *Mapk14* (or *p38α*), and the MAPK phosphatase *Dusp8* were all robust transcript heterogeneities in KP1 liver colonies (**Fig. 3C**). Clustering the 10-cell fluctuations of these genes indicated that elevated *Mapk14* levels co-occurred with reduced abundance of *Mapk8* and *Dusp8* (1 – *p* < 0.1 for co-enrichment; **Fig. 3G**). Signaling along these parallel MAPK effector pathways may be weighted differently among SCLC cells in the liver colony. Although no relevant receptors were detectably overdispersed in KP1 cells, we consistently detected *Il1rl1*, which is the receptor for Il33 of the interleukin 1 family. PNECs normally receive Il33 stimulation as an alarmin from ATII cells during lung injury or infection (55), but Il33 is also highly expressed in hepatocytes and liver sinusoids (33,56). The widespread single-cell adaptations downstream of Il33 support the hypothesis that SCLC cells redeploy native damage-response pathways in the liver microenvironment.

### Immunocompetency exaggerates stromal non-NE phenotypes in SCLC liver colonies

We built upon the results in athymic nude mice by repeating the liver colonization experiments in C57BL/6 x 129S F1 hybrid mice. Compared to KP1 colonies in athymic mice, the C57BL/6 x 129S F1 hybrid colonies had a higher proportion of CD3+ T cells as expected and a reduced proportion of F4/80+ macrophages along the colony margin (**Fig. 4A–D**), indicating different microenvironments. Stochastic profiling of the KP1 colony margins was performed in C57BL/6 x 129S F1 hybrid livers exactly as for athymic nude animals (**Fig. 3B**). Liver markers in the C57BL/6 x 129S F1 hybrid samples were as low and uncorrelated as in the nude samples (Supplementary Fig. S10). From 31 10-cell transcriptomic profiles, we robustly identified 1025 regulatory heterogeneities within KP1 cells colonized to an immunocompetent liver (**Fig. 4E** and Supplementary File S1).

**Figure 4.**
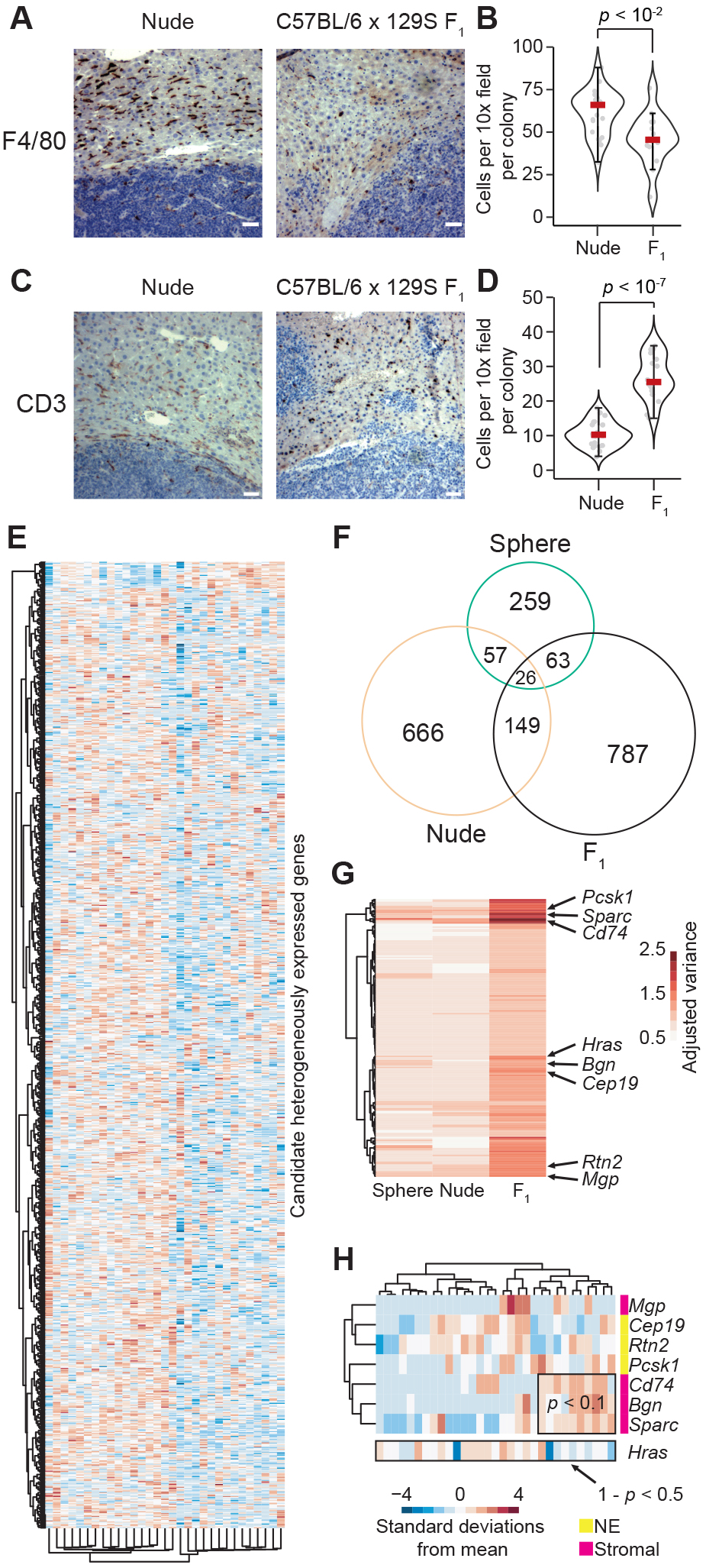
Stromal markers emerge heterogeneously when KP1 cells colonize immunocompetent liver. **A,** Immunohistochemistry of athymic nude (left) and C57BL/6 x 129S F1 hybrid (right) livers stained for the macrophage marker F4/80. **B,** Quantification of F4/80+ cells per 10x field is shown as the median of *n* = 23 nude colonies and *n* = 14 C57BL/6 x 129S F1 hybrid colonies. **C,** Immunohistochemistry of athymic nude (left) and C57BL/6 x 129S F1 hybrid (right) livers stained for the T cell marker, CD3. **D,** Quantification of CD3+ cells per 10x field is shown (right) as the median of *n* = 24 nude colonies and *n* = 23 C57BL/6 x 129S F1 hybrid colonies. **E,** Log-standardized 10-cell sampling fluctuations of robust candidate heterogeneities identified by stochastic profiling. **F,** Venn diagram comparing the heterogeneous transcripts identified in **Fig. 2B**, **3C**, and **E**. All two- and three-way intersections were significant (*p* < 0.001 by Monte-Carlo simulation). **G,** Regulatory heterogeneities with abrupt increases in abundance-adjusted variance in C57BL/6 x 129S F1 hybrid liver colonies. Stromal and neuroendocrine (NE) markers are highlighted. **H,** Log-standardized sampling fluctuations for the markers highlighted in **G.** Enriched or unenriched coexpression was evaluated by hypergeometric test of 10-cell observations above their respective log means. For **B** and **D**, differences were assessed by rank-sum test. Scale bar in **A** and **C** is 80 μm.

Gene set enrichment of the C57BL/6 x 129S F1 hybrid candidates reconstituted most of the hallmarks identified previously along with a moderate signature for hypoxia (Supplementary File S2). The hypoxia signature did not suggest preselection of cell subpopulations from the originating spheroid culture, as KP1 cells on the spheroid periphery showed the same range of Hif1α stabilization as cells in the spheroid core (Supplementary Fig. S11). In search of shared themes, we compared the KP1 candidate genes from the three biological contexts and found that all two- and three-way intersections were significant (*p* < 0.001 by Monte-Carlo simulation; **Fig. 4F**). This suggested that biological meaning might be embedded in the heterogeneity trends between groups. In lieu of fixed overdispersion thresholds (as in **Fig. 2A**), we next analyzed the adjusted variance as a continuous measure of predicted heterogeneity. Beginning with the 2007 transcripts predicted to be heterogeneously regulated in at least one context (**Fig. 4F**), we searched for genes with significant overdispersion increases in the immunocompetent setting (see Materials and Methods). We identified 202 transcripts meeting these criteria, which included multiple neuroendocrine markers (*Rtn2, Pcsk1, Cep19*), *Cd74*, and a new group of stromal transcripts (*Bgn, Sparc, Mgp;* **Fig. 4G**) (1). Mesenchymal transitions of SCLC cells can be driven by activated Kras (23), and we noticed that the dispersion of wildtype *Hras* increased alongside the stromal transcripts. However, when 10-cell fluctuations were clustered, we found that the co-occurrence of *Cd74–Bgn–Sparc* associated with a lack of elevated *Hras* abundance (**Fig. 4H**), excluding a straightforward EMT-like state change. The stromal marker *Mgp* was also uncoupled from *Cd74–Bgn–Sparc.* We conclude that immunocompetency drives a further diversification of SCLC toward stromal phenotypes in the setting of liver colonization.

Among the stromal transcripts, there was a particularly stark difference in *Cd74* dispersion between the two liver settings (**Fig. 4G**). We validated this finding by Cd74 immunofluorescence of separate GFP^+^ liver colonies among independent animals. Using automated image segmentation (see Materials and Methods), we detected GFP^+^Cd74^+^ cells in 52/54 = 96% of colonies imaged across both strains. Further, the variability in GFP^+^ Cd74^+^ content among colonies within a mouse was comparable to that among mice (*p* = 0.13 by random-effects two-way ANOVA after arcsine square-root transformation of percentages), enabling aggregate analysis. In athymic nude mice, we classified 3730/39593 = 9.4% of GFP^+^ cells as Cd74^+^, supporting the gain in regulatory state identified earlier by stochastic profiling (**Fig. 3D** and Supplementary Fig. S12A and S12B). In C57BL/6 x 129S F1 hybrid animals, the percentage increased significantly to 6894/26027 = 26% (*p* ~ 0 by binomial test; Supplementary Fig. S12C and S12D). Both settings required a two-state distribution to capture the Cd74 immunoreactivity (Supplementary Fig. S12B and S12D). This independent follow-up confirmed the accuracy of the stochastic-profiling results for *Cd74* across multiple in vivo settings, and the quantitative differences suggested a role for adaptive immune cells in driving the heterogeneity.

### Marker gene aberrations are partly retained in autochthonous SCLC tumors and metastases

The non-NE markers identified by stochastic profiling prompted a more-systematic evaluation of marker-gene status in 10-cell and bulk samples. For comparison, we used RNA-seq data from autochthonous tumors and metastases of *Rb1^F/F^Trp53^F/F^Rbl2^F/F^* mice administered AdCMV-Cre or adenoviral Cre driven the Calca promoter (AdCalca-Cre) (21). Curiously, for the ATII markers *Cd74* and *Lyz2*, the autochthonous samples indicated that abundance was higher in the primary tumor and reduced in the metastasis (**Fig. 5A** and **5B**, black squares vs. brown filled triangles). Similar results were obtained with the stromal markers, *Bgn* and *Sparc*, although the tumor-metastasis differences were less dramatic (**Fig. 5C** and **5D**). These observations are reconcilable with the 10-cell data if the spheroid observations are not taken as a proxy for the primary tumor. Rather, in vitro cultures reflect the SCLC states achievable from purely homotypic cell-cell interactions. Paracrine inputs from non-NE cells of the lung could just as feasibly drive SCLC reprogramming as non-NE cells of the liver.

**Figure 5.**
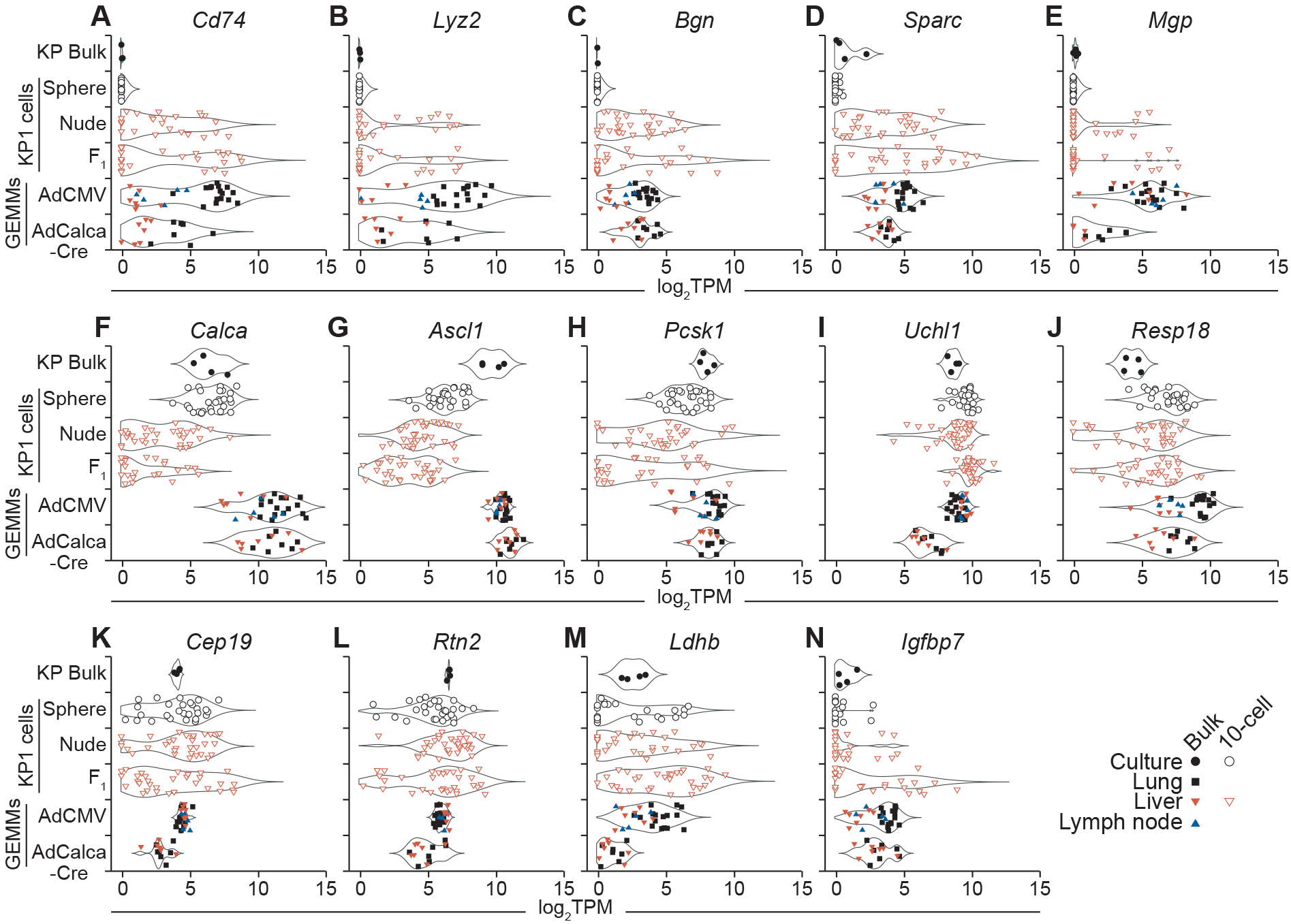
Comparison of 10cRNA-seq observations to bulk RNA-seq from primary SCLC tumors and metastases of SCLC GEMMs. **A** and **B,** Sporadic expression of the ATII markers *Cd74* (**A**) and *Lyz2* (**B**) upon liver colonization of KP1 cells compared to SCLC GEMM tumors and metastases. **C–E,** Abundance changes in the stromal markers *Bgn* (**C**), *Sparc* (**D**), and *Mgp* (**E**) in vivo. **F–J,** Reduced in vivo abundance of the neuroendocrine markers *Calca* (**F**), *Ascl1* (**G**), and *Pcsk1* (**H**), but not *Uchl1* (**I**) or *Resp18* (**J**). **K** and **L,** Context-dependent dispersion changes without abundance changes for the neuroendocrine markers *Cep19* (**K**) and *Rtn2* (**L**). **M,** The non-neuroendocrine marker *Ldhb* is a RHEG in KP1 cells. **N,** Heterogeneous regulation of the non-neuroendocrine marker *Igfbp7* in KP1 cells colonized to the liver of immunocompetent animals.

Interestingly, abundance of the stromal marker *Mgp* was quite different between the two autochthonous GEMMs (**Fig. 5E**). KP1 cells were isolated from an animal infected with AdCMV-Cre (24), and the lack of detectable *Mgp* in cultured KP1 cells suggests a liver-dependent induction. The sporadic increases in *Mgp* abundance observed upon liver colonization were consistent with the other stromal markers found to be high in AdCMV-Cre tumors and metastases. By contrast, *Mgp* abundance in AdCalca-Cre-derived samples was uniformly low. AdCalca-Cre has been speculated to target a more-differentiated subset of PNECs compared to AdCMV-Cre (21), even though *Calca* levels in the two tumor groups are roughly equivalent (**Fig. 5F**). We found that the gained *Mgp* expression of KP1 cells in vivo coincided with loss of endogenous *Calca* itself (**Fig. 5E** and **5F**). *Mgp* is an inhibitory morphogen for lung development (57) and its inducibility may mark the PNEC progenitor pool targeted by AdCMV-Cre.

In addition to *Calca*, other neuroendocrine markers (*Ascl1, Pcsk1*) declined substantially when KP1 cells were engrafted to the liver (**Fig. 5G** and **5H**). Yet, dedifferentiation appeared incomplete, as multiple neuroendocrine markers (*Uchl1, Resp18*) remained largely unchanged and at comparable abundance to autochthonous models (**Fig. 5I** and **5J**). Among the markers identified by a gradient of 10-cell dispersion (**Fig. 4G**), several showed no discernible change in median abundance (**Fig. 5K** and **5L**) and thus would be impossible to identify in bulk samples. One of the transcripts correlating strongly with non-NE markers in PNECs (*Ldhb*) (1) recurred as a candidate heterogeneity in all three KP1 settings (**Fig. 5M**). Lastly, we identified a characteristic non-NE marker (*Igfbp7*) (23) where both median abundance and dispersion increased specifically in immunocompetent livers (**Fig. 5N**). Such miscoordination of markers could occur if SCLC cells fragmented their transcriptional regulation upon encountering progressively more-diverse cellular microenvironments.

### Notch2 signaling is rapidly activated during KP1 cell dissociation

KP1 cells expressed multiple non-NE transcripts in the liver of C57BL/6 x 129S F1 hybrid mice (**Fig. 4G** and **4H**), and single-cell transcriptomics has associated non-NE changes with activation of the Notch pathway (1,12). Unexpectedly, despite measurable expression of *Notch2* in KP1 cells by 10cRNA-seq (3.4 ± 9.4 TPM), we almost never detected the Notch target gene *Hes1* in vivo (0 ± 0.1 TPM). Normal PNECs change to a non-NE state during tissue damage, which may be mimicked by the cell-dissociation steps required for conventional scRNA-seq (1). Notch-pathway activation of cell lines also reportedly occurs during routine passaging (58), prompting us to ask whether such artifacts could arise in KP1 cells. Notch1 is nearly absent in the line (less than 0.5 TPM for *Notch1* vs. 29 TPM for *Notch2* in bulk; GSE147358), and reliable activation-specific antibodies for Notch2 are not available. Therefore, we used an antibody recognizing an intracellular epitope of full-length Notch2 and its processed transmembrane (NTM) subunit, which is the precursor for pathway activation (59). Within five minutes of KP1 dissociation using either trypsin or accutase, we noted considerable decreases in total Notch2 protein (full-length + NTM; **Fig. 6A–C**). Furthermore, trypsin significantly increased the ratio of NTM-processed to full-length Notch2 (*p* < 0.01 by ANOVA; **Fig. 6D**), suggesting that trypsinized cells may be more primed to activate Notch signaling. One hour later—a typical delay for scRNA-seq—both dissociating enzymes led to induction of the Notch target genes, *Hes1* and *Hey1* (**Fig. 6E–G**). Our results support earlier speculation (1) that Notch activation in PNEC-like cells may be an artifact of the sample processing that precedes scRNA-seq but is avoided by 10cRNA-seq (25).

**Figure 6.**
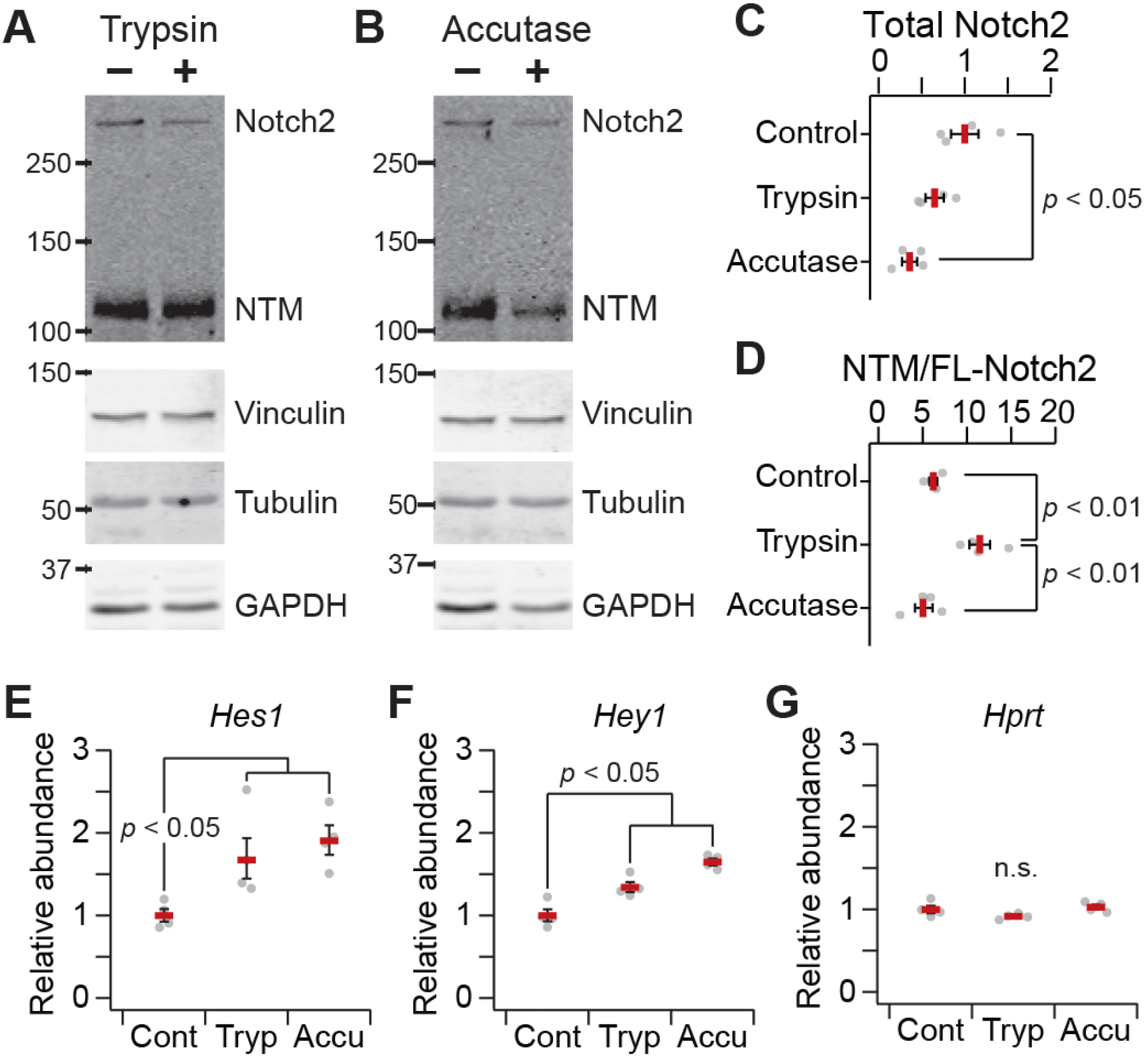
Cell-dissociation enzymes rapidly disrupt intracellular precursors of Notch2 signaling and induce Notch target genes in KP1 cells. **A** and **B,** Immunoblots of full-length Notch2 and the processed Notch transmembrane (NTM) subunit in KP1 cells after treatment of 0.05% trypsin or 1x accutase for five minutes. Vinculin, tubulin, and GAPDH were used as loading controls. **C,** Relative abundance of total Notch2 (full-length + NTM) for the indicated conditions. Data are normalized to control KP1 cells lysed without dissociation. **D,** Ratiometric abundance of NTM / full-length (FL) Notch2 for the indicated conditions. **E–G,** Quantitative PCR of the Notch target genes *Hes1* (**E**) and *Hey1* (**F**) and the housekeeping gene *Hprt* (**G**) in KP1 cells one hour after treatment of 0.05% trypsin or 1x accutase for five minutes. Samples were normalized to *Rpl30* and *Ppia* as loading controls. For **C** and **D**, data are shown as the mean ± s.e.m. from *n* = 4 independent biological samples. For **E–G**, data are shown as the geometric mean ± log-transformed s.e. from *n* = 4 independent biological samples. Differences in means were assessed by ANOVA with Tukey HSD post-hoc test (**C** and **D**) or log-transformed Student’s *t* test with Šidák correction (**E–G**).

### Human SCLCs are merged or stratified by different classes of KP1 RHEGs

We returned to two statistically significant overlaps from the three studies in KP1 cells (**Fig. 4F**). The three-way intersection of 26 transcripts defined a core group of RHEGs, which we viewed as a set of cell-autonomous heterogeneities intrinsic to KP1 cells and perhaps SCLCs more generally. We tested this concept by identifying the human orthologs of the KP1 core RHEG set and clustering our data alongside bulk RNA-seq profiles from 79 cases of SCLC in humans (32). The standardized fluctuations of the core RHEGs in human samples were largely indistinguishable from the KP1 observations, with most sample co-clusters containing mouse and human data (**Fig. 7A**). Moreover, when the pairwise correlations of core RHEGs were organized hierarchically, it was difficult to discern any strongly linked groups of observations (**Fig. 7B**). This would be expected if core RHEGs were broadly but independently “active” (induced heterogeneously). Accordingly, we found very little evidence of coordination outside a small row cluster of genes involved in biological processes that were largely unrelated—cell cycle-dependent ubiquitination (*CCNF*), carbonyl stress (*HAGH*), splicing (*SNRNP200*), calcium homeostasis (*CHERP*), and DNA methylation (*MBD1*) (**Fig. 7A**). Although the existence of core RHEGs in mammalian SCLCs awaits direct testing in human samples, the analysis here provides a GEMM-informed set of targets worth examining further.

**Figure 7.**
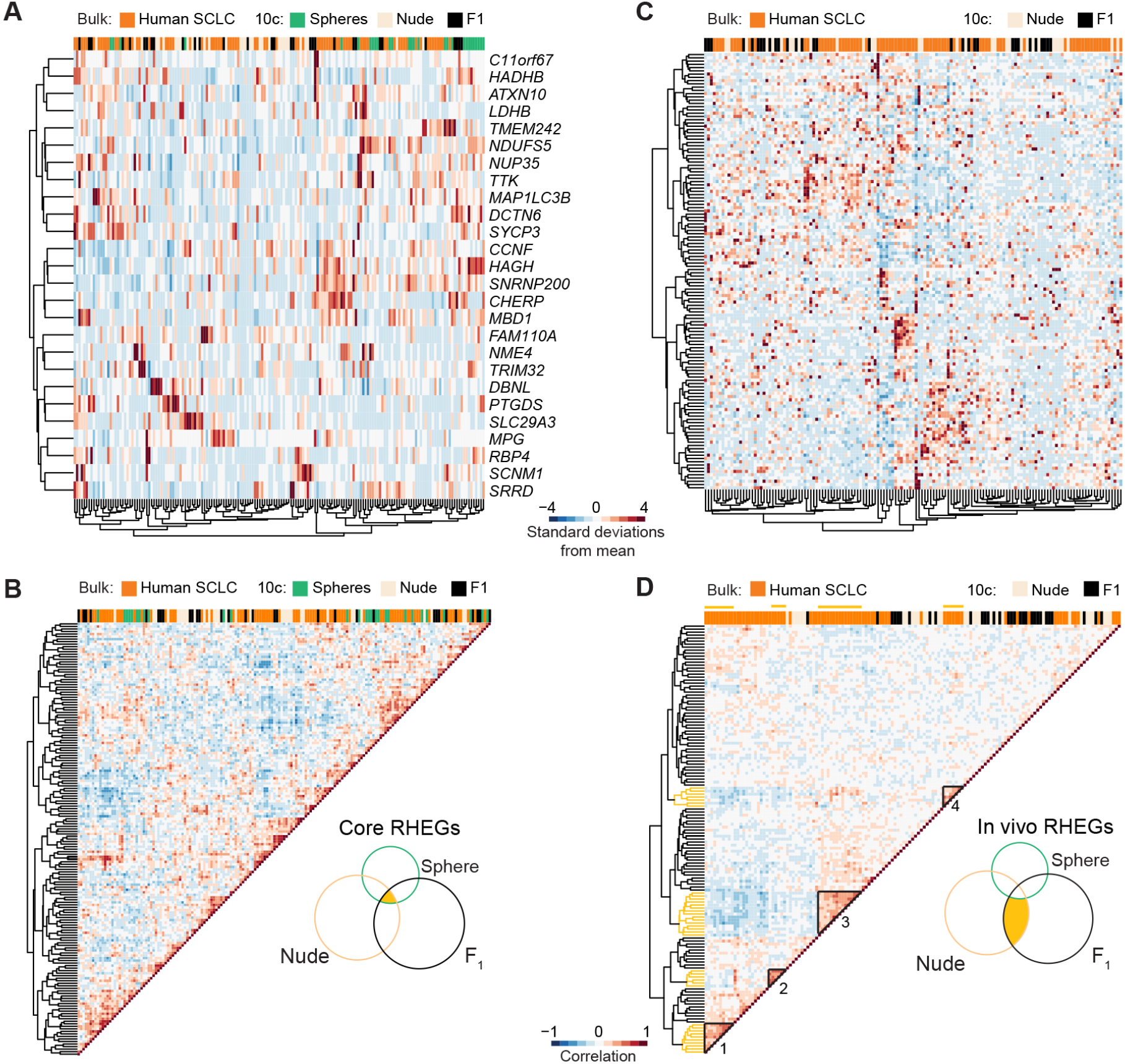
Orthologous RHEG fluctuations in primary human SCLCs. **A,** Core RHEGs and their orthologs intermix KP1 10-cell observations and bulk RNA-seq data from human cases of SCLC (32). **B,** Pearson correlation matrix for core RHEGs clustered hierarchically. The Venn diagram intersection for core RHEGs is highlighted from **Fig. 4F. C,** In vivo RHEGs and their orthologs do not merge KP1 and human observations but identify subgroups of clinical SCLCs. **D,** Pearson correlation matrix for in vivo RHEGs clustered hierarchically. Groups of covarying human SCLC cases are indicated in black triangles and yellow margins and numbered as hierarchical clusters in Supplementary Fig. S13. The Venn diagram intersection for core RHEGs is highlighted from **Fig. 4F**. Murine data and human data were standardized separately by z-score before clustering or correlation.

The second overlap of interest was the two-way intersection of 149 genes that emerged as candidate heterogeneities in both settings of liver colonization (**Fig. 4F**). We defined these in vivo RHEGs as reflecting the SCLC regulatory heterogeneity triggered by heterotypic cell-cell interactions in the microenvironment. In contrast to core RHEGs, we expected different activation patterns of in vivo RHEGs in the liver versus the lung, and even among different SCLC subtypes or primary-tumor sites in the lung. We extracted human orthologs of the in vivo RHEGs and clustered the KP1 observations together with the human SCLCs (**Fig. 7C**). There was far less intermixing between human and KP1 samples, consistent with the different heterotypic interactions anticipated between primary and metastatic sites. The pairwise correlation structure of in vivo RHEGs was also qualitatively distinct, with multiple groups of covariates comprised entirely of human SCLCs (**Fig. 7D**). We found by consensus clustering (39) that three of the four human groups were robust (Supplementary Fig. S13A). Importantly, these clusters each contained mixtures of various SCLC subtypes based on the relative abundance of key transcription factors (18) (Supplementary Fig. S13B). Clinical outcomes were available for only 48 sequenced SCLC cases (32). Nevertheless, those in the first in vivo RHEG cluster showed significantly longer survival compared to cases that did not cluster (*p* < 0.01 by log-rank test; Supplementary Fig. S13C). The KP1 observations in the liver reflect one SCLC subtype and do not precisely capture human variation in the lung. However, the in vivo RHEG set derived from those observations may stratify clinical cases by differences in tumor ecosystem and SCLC adaptation.

## Discussion

PNECs are a particularly versatile cell type (1), and it is perhaps unsurprising that derivative SCLC cells show the deranged plasticity reported here. Less obvious is whether dispersed SCLC states are engaged hierarchically or chaotically—our work with a representative GEMM-derived SCLC line argues for the former. Cell-autonomous regulatory heterogeneities expand qualitatively in vivo through heterotypic cell-cell interactions absent from in vitro culture. The documented cell-state changes upon liver colonization could simply reflect the injury-like state of tumors and metastases (6). Alternatively, the reprogramming events could provide trophic support to the cellular ecosystem (2,12). The candidate heterogeneities identified by stochastic profiling and 10cRNA-seq create a resource to guide future functional studies that perturb specific emergent heterogeneities in vivo.

The KP1 results with Notch2 reinforce that SCLC cells are very sensitive to juxtacrine inputs (12). SCLC tumorsphere growth in vitro elicits its own cell-to-cell heterogeneities, which have some commonalities with spheroids of MCF10A-5E basal-like breast cells, a distant epithelial cell type. Intrinsic to spheroid culture are subclonal reorganization and competition, two processes important for primary tumor initiation and the end stages of metastatic colonization. Cell crowding and sequestration alter lipid metabolism (60,61), which could explain the catabolic and anabolic lipid enzymes identified within the spheroid RHEG set. The notion of spheroid RHEGs may generalize to clonogenic soft-agar assays of anchorage-independent growth, which remain widely used as surrogates for tumorigenicity.

We recognize multiple limitations in the liver-colonization experiments. Intravenous delivery of tumor cells to a metastatic site is not the same as spontaneous metastases, which must disseminate from a primary tumor. Athymic nude and C57BL/6 x 129S F1 hybrid animals are very different from one another, and the genetic background may have altered the liver microenvironment even before colonization because of myeloid–lymphoid imbalances (52). KP1 cells are predominantly mixed from C57BL/6 and 129S strains, but we cannot exclude the possibility of immunogenic polymorphisms from other strains. Further, it is possible that the EGFP transgene added to KP1 cells exacerbates cell-mediated immunity in the C57BL/6 x 129S F1 hybrid context. We do not claim that liver colonization of C57BL/6 x 129S F1 hybrids captures authentic anti-SCLC immunity. For this study, it provided a context to evaluate how SCLC variability was boosted upon lymphocyte homing to the liver.

The candidate regulatory heterogeneities identified in KP1 liver colonies reflect several of the deprogramming and reprogramming events recently described in PNECs (1). In addition, they suggest routes of paracrine communication that are equally realistic for the lung as for the liver. From this perspective, the stratification of primary human SCLCs by in vivo RHEGs is intriguing, especially given the association of some RHEG clusters with patient survival. SCLCs usually initiate in the bronchi, but there are differences in cell composition at different depths of the lung (62) as well as lobular biases in the primary sites typical for SCLC (63). Different SCLC subtypes (18) might arise in similar microenvironments, yielding the mixed-subtype clusters identified here. The stromal heterogeneities induced by the immunocompetent setting may also relate to fibrotic lung diseases, where PNECs hyperplasia is known to occur (64). The genome of SCLCs is known to be highly mutated (13,14), but our study indicates that cell-fate variability arises on a much faster time scale in vivo. Primary tumors with coordinated RHEG-like signatures may be naturally trapped in a state of perpetual adaptation that ultimately slows progression and improves outcome. SCLC may be particularly amenable to therapeutic regimens that force transformed PNECs to adapt to constitutively changing environments (65).

## Acknowledgments

We thank Emily Farber and Suna Onengut-Gumuscu at the UVA Center for Public Health Genomics for RNA-seq library preparation and sequencing, Craig Rumpel and Patcharin Pramoonjago at the UVA Biorepository and Tissue Research Facility for LCM maintenance and histology services, UVA Research Computing for high-performance computing access and consulting, Henry Pritchard for assistance with NCBI GEO deposition, and Cheryl Borgman for critical evaluation of the manuscript. This work was supported by the National Institutes of Health #R01-CA194470 (K.A.J.), #U01-CA215794 (K.A.J.), #R01-CA194461 (K.S.P.), and #U01-CA224293 (K.S.P.), the David & Lucile Packard Foundation #2009-34710 (K.A.J.), a UVA Cancer Center support grant #P30-CA044579, the UVA Medical Scientist Training Program (S.S.), a Wagner Fellowship (S.S.), and a Harrison Undergraduate Research Award (D.L.S.).

