## Supplementary Files and Tables for "Fragmentation of Small-cell Lung Cancer Regulatory States in Heterotypic Microenvironments"

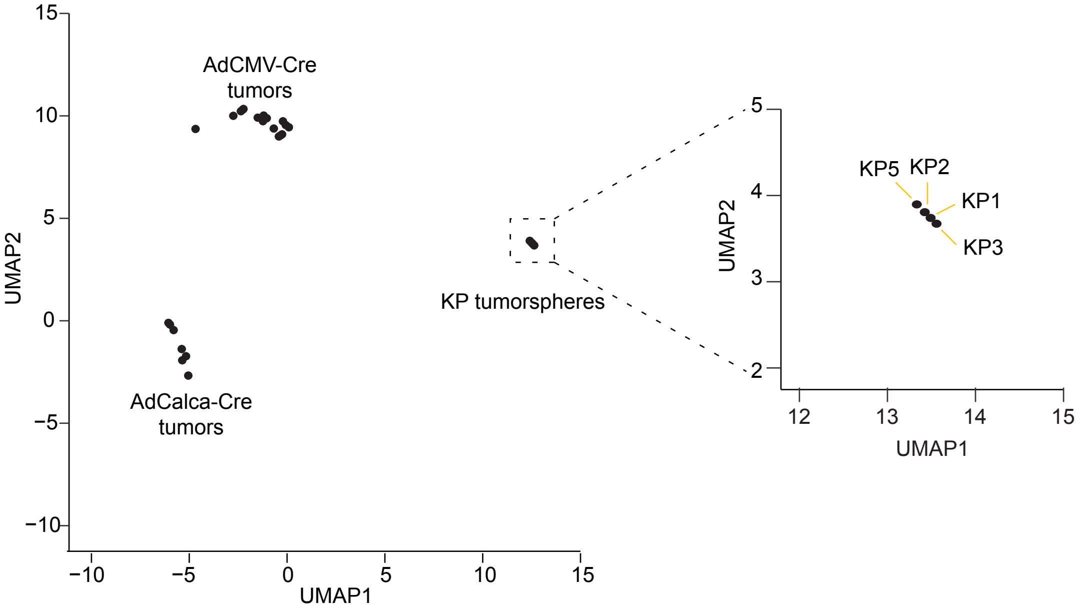


**Supplementary Figure S1.** KP1 cells are representative of SCLC lines derived from *Rb1*$^{F/F}$*Trp53*$^{F/F}$ mice administered CMV-driven adenoviral Cre (AdCMV-Cre). Autochthonous AdCMV-Cre-initiated and AdCalca-Cre-initiated tumors from *Rb1*$^{F/F}$*Trp53*$^{F/F}$*Rbl2*$^{F/F}$ mice (1) are shown for comparison. Genotypes for the KP SCLC lines are: KP1, *Rb1*$^{\Delta/\Delta}$*Trp53*$^{\Delta/\Delta}$ (2); KP2, *Rb1*$^{\Delta/\Delta}$*Trp53*$^{\Delta/\Delta}$*Ptch1*$^{+/Lacz}$ (2); KP3, *Rb1*$^{\Delta/\Delta}$*Trp53*$^{\Delta/\Delta}$*Axin2*$^{+/Lacz}$ (3); KP5, *Rb1*$^{\Delta/\Delta}$*Trp53*$^{\Delta/\Delta}$*Tg*$^{BAT\text{-}lacZ}$ (4).

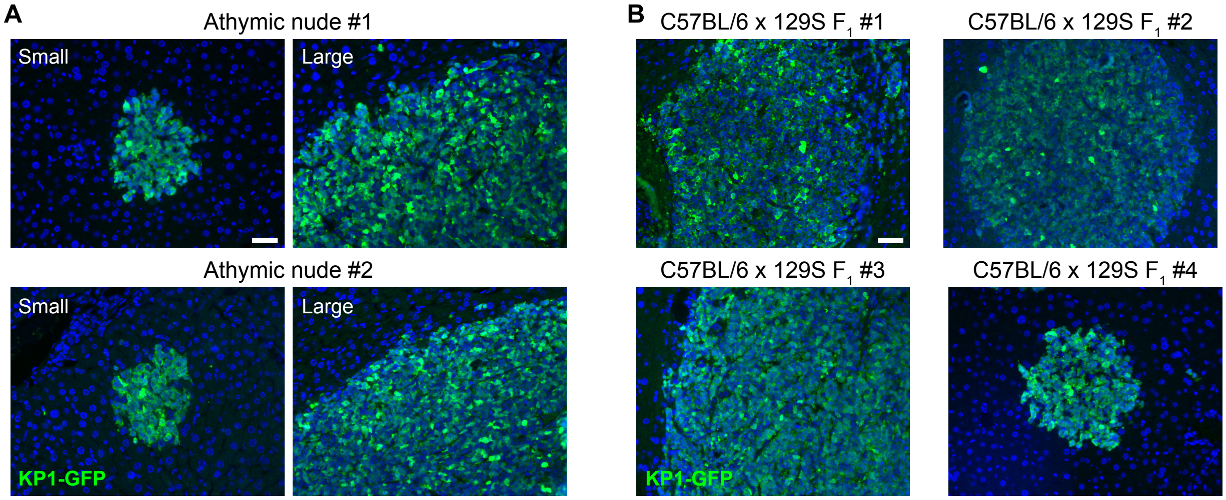


**Supplementary Figure S2.** Representative images of $GFP^+$ liver colonies upon intravenous administration of KP1-GFP cells. **A,** Small and large colonies in athymic nude animals. **B,** Various colonies in C57BL/6 x 129S $F_1$ hybrid animals. Scale bar is 40 μm.

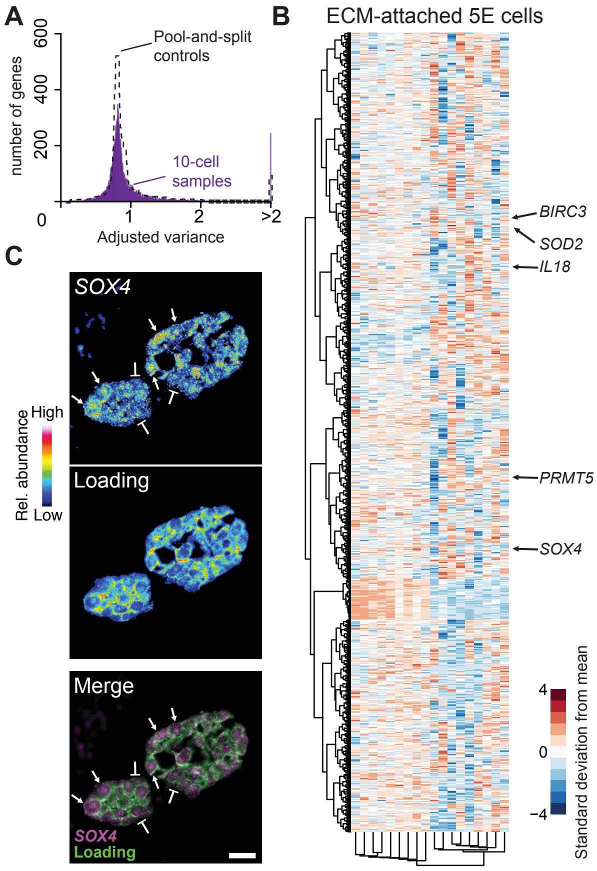


**Supplementary Figure S3.** Stochastic profiling of transcriptional regulatory heterogeneities in MCF10A-5E spheroids by 10cRNA-seq. **A,** Overdispersion plot showing 10-cell sample distribution for MCF10A-5E spheroids (purple) overlaid on pool-and-split controls of 10-cell equivalents of KP1 spheroids (dashed). **B,** Clustergram of the 1129 transcripts identified as candidate heterogeneities within MCF10A-5E spheroids. Data were log transformed before standardization. Indicated transcripts were independently validated as heterogeneous by RNA FISH (5,6). **C,** RNA FISH validation of *SOX4*, a gene candidate identified by 10cRNA-seq stochastic profiling. Pseudocolor image for *SOX4* is shown above a loading control hybridization comprised of *GAPDH*, *HINT1*, and *PRDX6* (7). Scale bar is 20 μm.

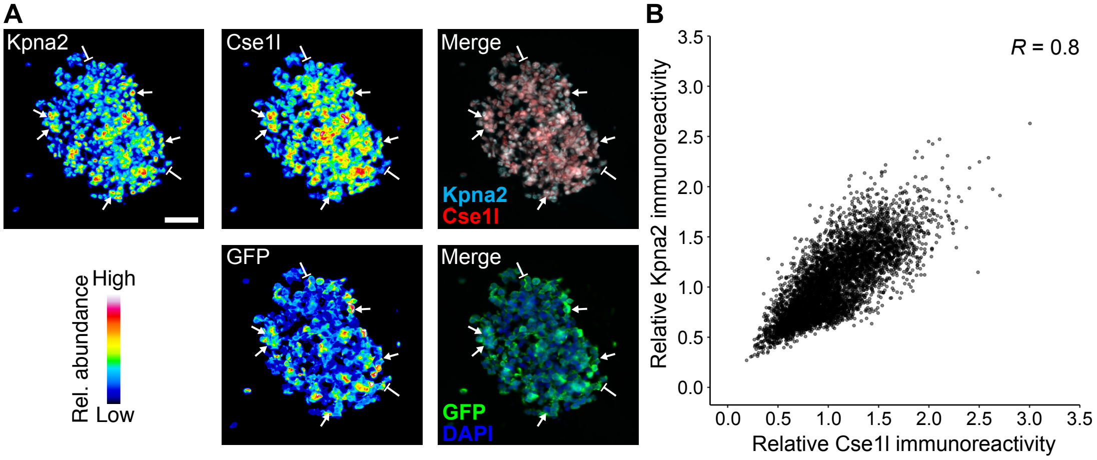


**Supplementary Figure S4.** Coregulation of Kpna2 and Cse1l in KP1 tumorspheres confirmed by quantitative immunofluorescence. **A,** Three-color immunostaining of Kpna2 (cyan), Cse1l (red), and GFP (green). Pseudocolored single-channel images are shown alongside the indicated merges. Scale bar is 40 μm. **B,** Segmented median-normalized Kpna2–Cse1l immunoreactivity in $n$ = 5741 cells from KP1-GFP spheroids. The Pearson correlation ($R$) is shown.

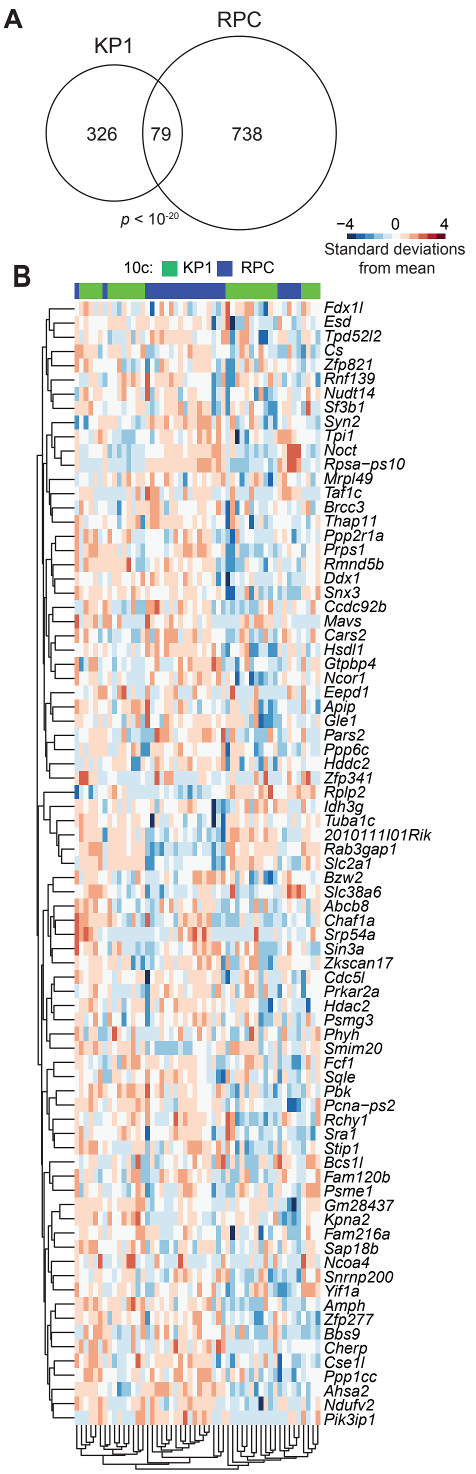


**Supplementary Figure S5.** Stochastic profiling of transcriptional regulatory heterogeneities in *Chga-GFP*;$Rb^{\Delta/\Delta}$;$p53^{\Delta/\Delta}$;$p130^{\Delta/+}$;$Crebbp^{-/-}$ (RPC) spheroids by 10cRNA-seq. **A,** Venn diagram of candidates between KP1 spheroids and RPC spheroids. Significance of the intersection was assessed by hypergeometric test with 12,612 total detectable transcripts in KP1 cells and 16,033 total detectable transcripts in RPC cells. **B,** Clustergram of the SCLC-spheroid RHEGs. Data were log transformed before standardization.

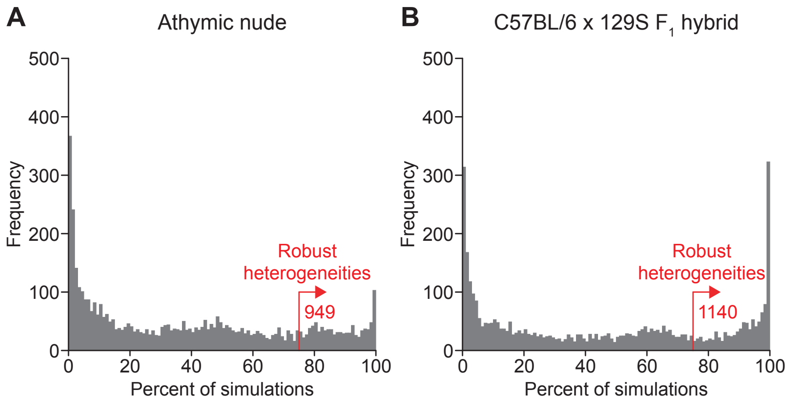


**Supplementary Figure S6.** Subsampling identifies robust transcriptional regulatory heterogeneities within KP1 liver colonies. **A,** Subsampled dispersion analysis of 33 10-cell observations of KP1 cells colonized to the liver of athymic nude mice. **B,** Subsampled dispersion analysis of 31 10-cell observations of KP1 cells colonized to the liver of immunocompetent C57BL/6 x 129S $F_1$ hybrid mice. Datasets were randomly downsampled to 28 10-cell observations and analyzed for overdispersion as described (8), and candidates appearing in >75% of subsampling runs were considered robust heterogeneities.

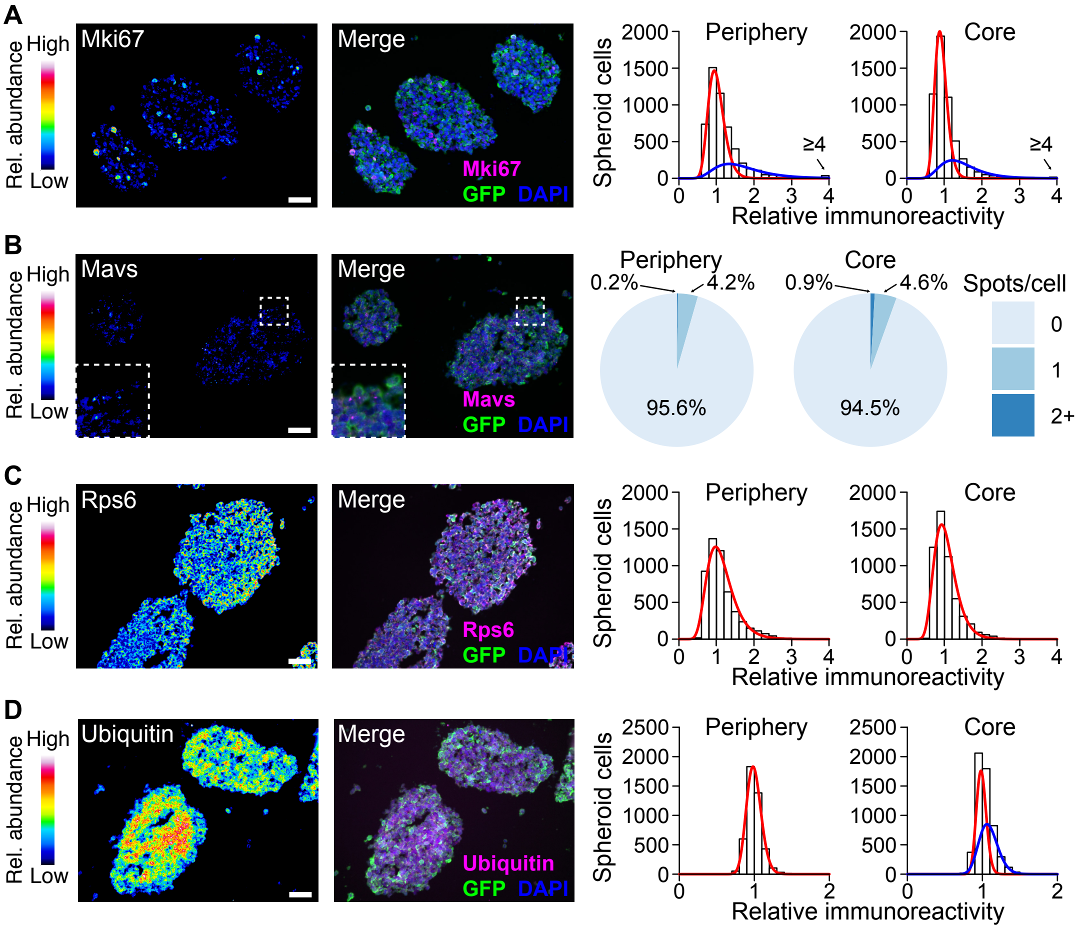


**Supplementary Figure S7.** Heterogeneous regulation of Mki67 and Mavs is comparable at the spheroid periphery and in the core. **A** and **B**, Two-color immunostaining of candidate heterogeneities Mki67 (**A**) or Mavs (**B**) and GFP. **C** and **D,** Two-color immunostaining of housekeeping protein Rps6 (**C**) and Ubc as total ubiquitin (**D**). Pseudocolored single-channel images are shown (left) alongside the indicated merge (center). Scale bar is 40 μm. The segmented immunoreactivity (right) was split based on spheroid location and fit to a one-state (red) or two-state (red and blue) lognormal distribution with model complexity assessed by *F* test, except for Mavs where immunoreactive spots (**B**, inset) were counted and summarized.

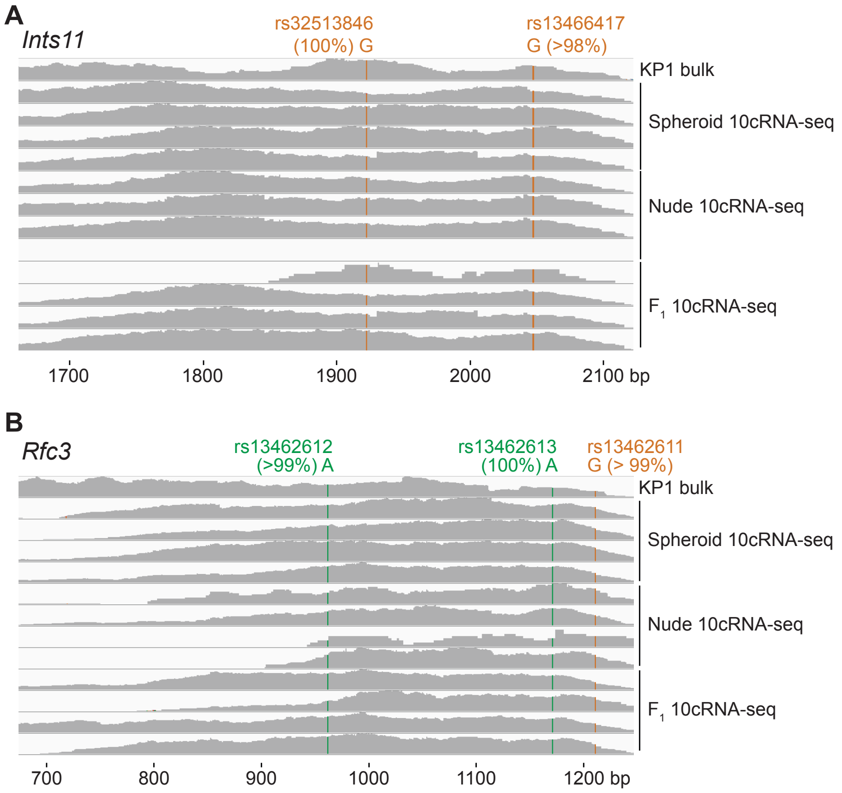


**Supplementary Fig. S8.** 10cRNA-seq transcriptomes from the liver colonies of athymic nude and C57BL/6 x 129S $F_1$ hybrid animals exhibit the single-nucleotide polymorphisms (SNPs) of KP1 cells exclusively. **A** and **B**, Bulk and 10cRNA-seq alignments of KP1 samples to the 3' region of *Ints11* (**A**) and *Rfc3* (**B**) in the C57BL/6 reference transcriptome. (*Ints11* was not detected in one athymic nude sample.) Homozygous SNPs deviating from the C57BL/6 transcriptome are highlighted along with an estimated allele frequency. The rs13462613 and rs13462611 SNPs in *Rfc3* are in linkage disequilibrium, and the AG block is absent from both C57BL/6 and 129S strains.

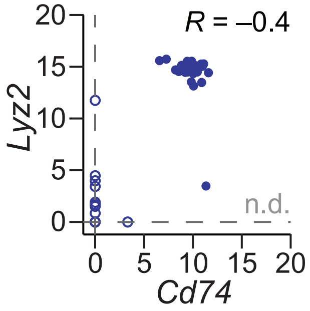


**Supplementary Figure S9.** *Cd74* and *Lyz2* are anti-correlated in single PNEC-derived non-NE cells. scRNA-seq reads of *Lyz2* and *Cd74* are shown for non-NE cells (9). n.d., not detected. Significance of the Pearson correlation (*R*) was tested after Fisher Z transformation (one-sided $p < 0.05$).

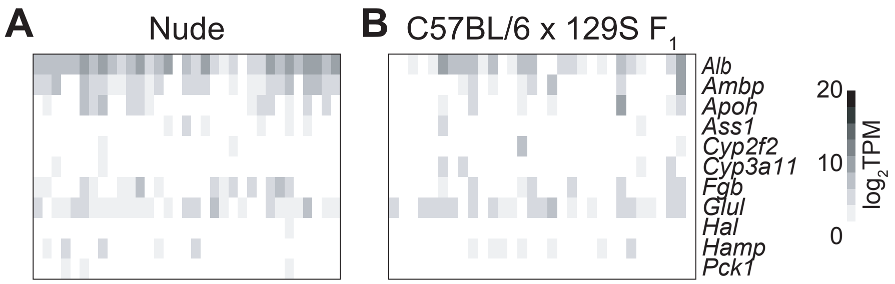


**Supplementary Figure S10.** Liver contamination in KP1 samples from C57BL/6 x 129S $F_1$ hybrid liver colonies is low and uncorrelated as in athymic nude liver colonies. **A,** Relative abundance of liver markers in 10-cell samples from nude liver colonies, reprinted from **Fig. 3C** for comparison. **B,** Relative abundance of liver markers in 10-cell samples from C57BL/6 x 129S $F_1$ hybrid liver colonies, column clustered as in **Fig. 4E**.

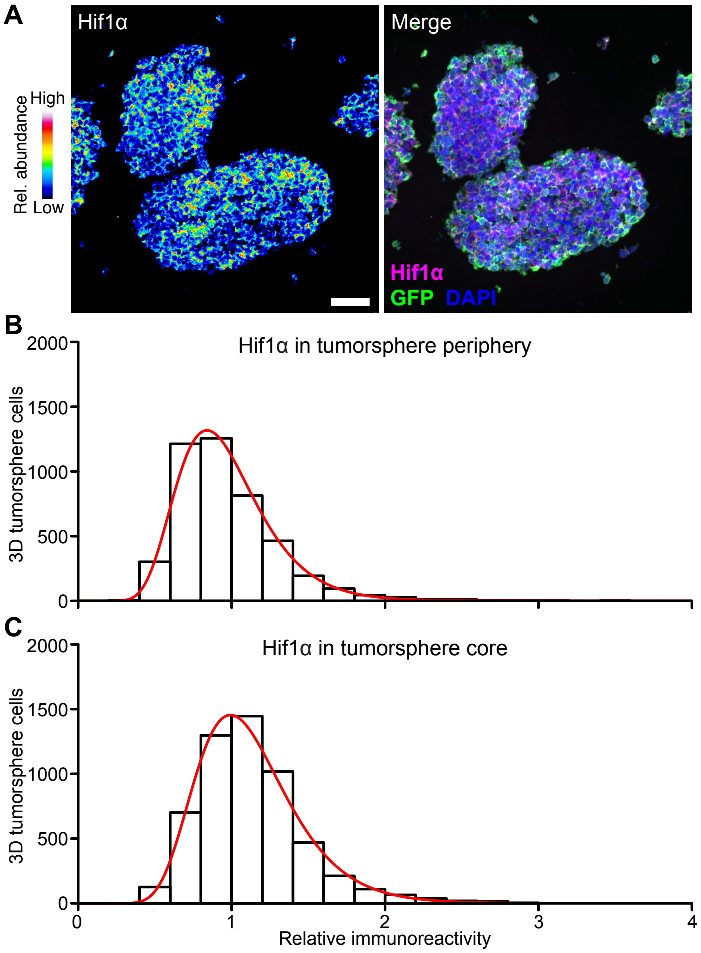


**Supplementary Figure S11.** Hif1α is not appreciably stabilized in the KP1 spheroid core or at the periphery. **A,** Two-color immunostaining of Hif1α (magenta) and GFP (green). A pseudocolored single-channel image is shown alongside the indicated merge. Scale bar is 40 μm. **B,** Segmented Hif1α immunoreactivity in $n$ = 9965 cells from KP1-GFP spheroids at the indicated locations.

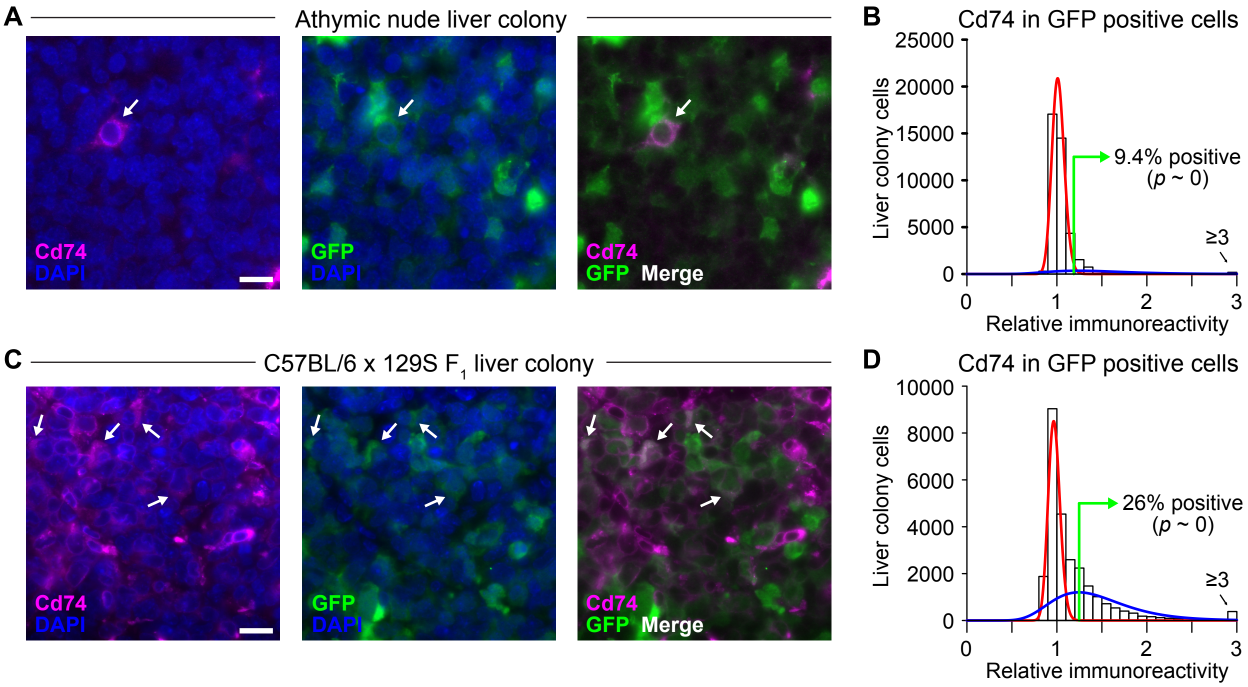


**Supplementary Figure S12.** Progressive gain in Cd74 expression among liver-colonized KP1-GFP cells confirmed by quantitative immunofluorescence. **A–D,** Two-color immunostaining of Cd74 (magenta) and GFP (green) for liver colonies in athymic nude mice (**A**) or C57BL/6 x 129S $F_1$ hybrid mice (**C**). Scale bar is 40 μm. Quantification of Cd74 immunoreactivity in segmented GFP+ cells across $n$ = 23 (athymic nude, **B**) or 31 (C57BL/6 x 129S $F_1$ hybrid, **D**) liver colonies. Significance of the positive subpopulation was assessed by binomial test with a 1% background probability.

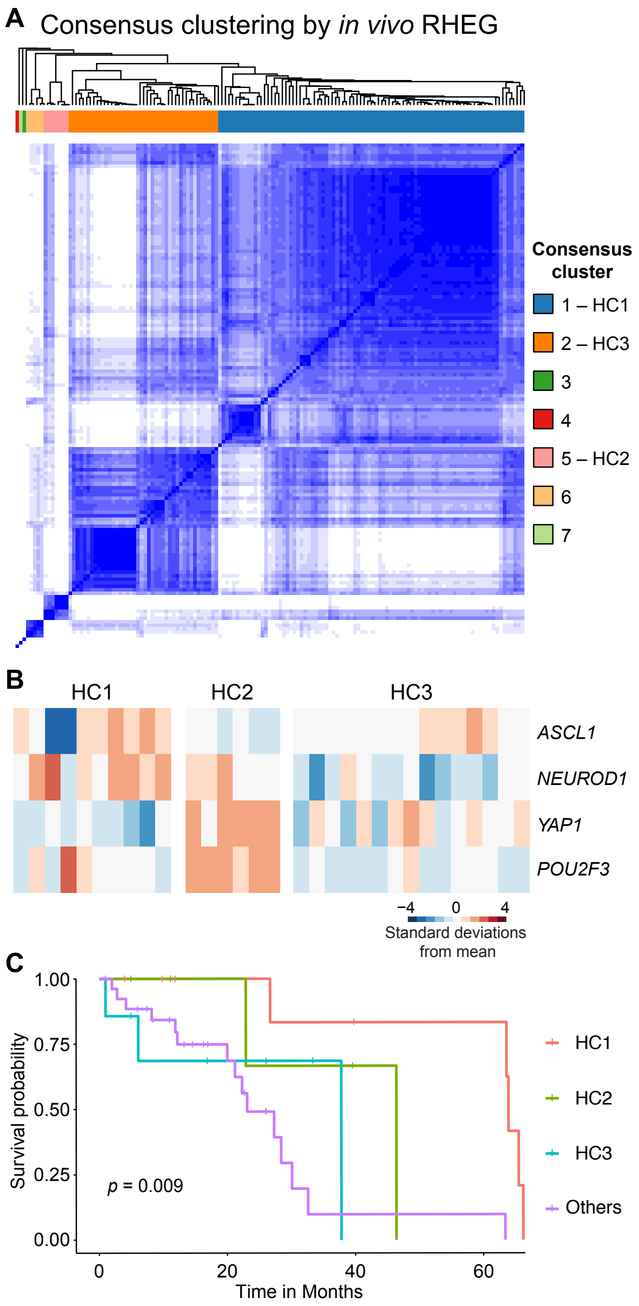


**Supplementary Figure S13.** Clinical mapping of in vivo RHEG clusters. **A,** Hierarchical clusters (HC) 1–3 (numbered as in **Fig. 7D**) are robust by consensus clustering. **B,** In vivo RHEG clusters of human SCLC are not entirely explained by known SCLC subtypes. The SCLC-A, SCLC-N, SCLC-Y, and SCLC-P subtypes are based on the above transcription factor abundances (10). **C,** Kaplan-Meier survival analysis of patients with clinical outcome (11).

**Supplementary Table ST1.** List of qPCR primers used in this study.

| Transcript | Forward primer | Reverse primer |
|---|---|---|
| *Gapdh* | GGCATTGCTCTCAATGACAA | GCCTCTCTTGCTCAGTGTCC |
| *Rpl30* | ATCCTTGCCAACAACTGTCC | CACCTGGGTCAATGATAGCC |
| *Alb* | CTGAACCGTGTGTGTCTGCT | TGGAAGGTGAAGGTCTCAGC |
| *Fgb* | ACTACTGGGGTGGCCTTTAC | ATGAAAGTCTGTGCTTGGGG |
| *Cyp3a11* | TGTACTGAATCTTTAACCAGGCA | TTGTTCTAAAGGTTGTGCCACG |
| *Hprt* | ATAGGCCAGCCTACCCTCTG | TCAATCACATCTTTCTCTCCTGA |
| *Ppia* | CACAAACGGTTCCCAGTTTT | AGCTGTCCACAGTCGGAAAT |
| *Hes1* | AAAGTCCCTAGCCCACCTCT | AAGGCGCAATCCAATATGAACA |
| *Hey1* | GCTTGGTGCCTGTGAAACAC | CCTAGTTCAGAGGAGCTGTTCA |

**Supplementary File S1.** List of candidate regulatory heterogeneities, adjusted dispersion values, and range of TPM values from stochastic profiling of 10-cell samples and split-pool controls. Individual worksheets are for SCLC spheroids of KP1-GFP cells (Sheet 1) or *Chga-GFP;Rb*$^{\Delta/\Delta}$*;p53*$^{\Delta/\Delta}$*;p130*$^{\Delta/+}$*;Crebbp*$^{-/-}$ (RPC) cells (Sheet 2) and KP1-GFP liver colonies in athymic nude (Sheet 3) or C57BL/6 x 129S $F_1$ hybrid (Sheet 4) animals.

**Supplementary File S2.** Hallmark gene set enrichments for transcripts with human orthologs in MSigDB. Individual worksheets are for SCLC spheroids of KP1-GFP cells (Sheet 1) or *Chga-GFP;Rb*$^{\Delta/\Delta}$*;p53*$^{\Delta/\Delta}$*;p130*$^{\Delta/+}$*;Crebbp*$^{-/-}$ (RPC) cells (Sheet 2) and KP1-GFP liver colonies in athymic nude (Sheet 3) or C57BL/6 x 129S $F_1$ hybrid (Sheet 4) animals.
